# Iron transport pathways in the human malaria parasite *Plasmodium falciparum* revealed by RNA-sequencing

**DOI:** 10.1101/2024.04.18.590068

**Authors:** Juliane Wunderlich, Vadim Kotov, Lasse Votborg-Novél, Christina Ntalla, Maria Geffken, Sven Peine, Silvia Portugal, Jan Strauss

## Abstract

Host iron deficiency is protective against severe malaria as the human malaria parasite *Plasmodium falciparum* depends on bioavailable iron from its host to proliferate. The essential pathways of iron acquisition, storage, export, and detoxification in the parasite differ from those in humans, as orthologs of the mammalian transferrin receptor, ferritin, or ferroportin, and a functional heme oxygenase are absent in *P. falciparum*. Thus, the proteins involved in these processes may be excellent targets for therapeutic development, yet remain largely unknown. Here, we show that parasites cultured in erythrocytes from an iron-deficient donor displayed significantly reduced growth rates compared to those grown in red blood cells from healthy controls. Sequencing of parasite RNA revealed diminished expression of genes involved in overall metabolism, hemoglobin digestion, and metabolite transport under low-iron versus control conditions. Supplementation with hepcidin, a specific ferroportin inhibitor, resulted in increased labile iron levels in erythrocytes, enhanced parasite replication, and transcriptional upregulation of genes responsible for merozoite motility and host cell invasion. Through endogenous GFP tagging of differentially expressed putative transporter genes followed by confocal live-cell imaging, proliferation assays with knockout and knockdown lines, and protein structure predictions, we identified six proteins that are likely required for ferrous iron transport in *P. falciparum*. Of these, we localized *Pf*VIT and *Pf*ZIPCO to cytoplasmic vesicles, *Pf*MRS3 to the mitochondrion, and the novel putative iron transporter *Pf*E140 to the plasma membrane for the first time in *P. falciparum*. *Pf*NRAMP/*Pf*DMT1 and *Pf*CRT were previously reported to efflux Fe^2+^ from the digestive vacuole. Our data support a new model for parasite iron homeostasis, in which *Pf*E140 is involved in iron uptake across the plasma membrane, *Pf*MRS3 ensures non-redundant Fe^2+^ supply to the mitochondrion as the main site of iron utilization, *Pf*VIT transports excess iron into cytoplasmic vesicles, and *Pf*ZIPCO exports Fe^2+^ from these organelles in case of iron scarcity. These results provide new insights into the parasite’s response to differential iron availability in its environment and into the mechanisms of iron transport in *P. falciparum* as promising candidate targets for future antimalarial drugs.

## INTRODUCTION

Iron is an essential micronutrient for all living organisms and has been associated with virulence of many pathogens. Iron abundance increases the replication of human immunodeficiency virus (HIV) (1) and *Mycobacterium tuberculosis* (2), and promotes biofilm formation in *Pseudomonas aeruginosa* (3). A “fight for iron” has been described between bacteria and the human host in the gastrointestinal tract (4), where the metal skews the composition of the gut microbiome by facilitating the growth of enteropathogenic *Escherichia coli* and *Salmonella* (5). Similarly, cancer cells require more iron compared to healthy cells (6) and higher ferritin levels in individuals diagnosed with COVID-19 were associated with increased disease severity and lethality (7).

Host iron deficiency is known to be protective against severe malaria (8–11) and iron chelators have cytocidal effects on the human malaria parasite *Plasmodium falciparum* (12). This obligate intracellular parasite depends on bioavailable iron for its proliferation and relies entirely on the host to meet its nutrient requirements (13). Furthermore, *P. falciparum* senses environmental fluctuations (14–16) and modulates its virulence in response (16). While iron is crucial for DNA replication and repair, mitochondrial electron transport, and redox regulation, it becomes toxic when in excess, as it is a source of damaging reactive oxygen species (17). Importantly for therapeutic development, the mechanisms of iron acquisition, storage, detoxification, and export in the parasite are different from those in humans, as orthologs of the mammalian transferrin receptor, ferritin, or ferroportin, and a functional heme oxygenase are absent in *Plasmodium* (18).

While human blood plasma contains between 10 and 30 µM total iron and an erythrocyte carries approximately 20 mM Fe (19), only 3 µM labile iron is present in the cytosol of uninfected red blood cells, and 1.6 µM in *P. falciparum*-infected ones (20). An estimated total iron concentration of 500 mM (21) is reached within the parasite’s digestive vacuole (DV), where iron-containing hemoglobin (Hb) is digested and the released heme is detoxified by biocrystallization into hemozoin (22). However, *P. falciparum* cannot access this iron source and is thought to acquire bioavailable Fe^2+^ from the host cell (23). Over-elevated ferrous iron levels likely compromise the integrity of the DV membrane and cytosolic iron also needs to be regulated to prevent oxidative stress (18). Iron detoxification in the parasite can be achieved by translocating the metal ion into dynamic intracellular Fe^2+^ stores, which may include acidocalcisomes – cytoplasmic vesicles that contain high concentrations of phosphate, calcium, iron, and zinc (24). In contrast to *Trypanosoma brucei* (24), no transport proteins have yet been experimentally shown to localize to the acidocalcisome membrane in *P. falciparum* (25, 26). Like the DV, these organelles are thought to be acidified by the plant-like V-ATPase and their low internal pH may fuel secondary active transport processes (22, 27).

*P. falciparum* encodes approximately 200 transmembrane or membrane-associated transport proteins (channels, pores, carriers, and pumps), many of which are essential for parasite growth and lack human homologs (28). For instance, the vacuolar iron transporter *Pf*VIT (PF3D7_1223700), an ortholog of *Arabidopsis thaliana* VIT1 (expect value (E) = 5 x 10^-29^, 30.5% identity, 87% coverage, as determined by position-specific iterated BLAST (29)), is a Fe^2+^/H^+^ exchanger that plays a role in iron detoxification (30–32). While its orthologs were localized to the endoplasmic reticulum in *Plasmodium berghei* (30) and to the vacuolar compartment in *Toxoplasma gondii* (33), the subcellular localization in *P. falciparum* had not been investigated experimentally prior to this study. Similarly, the Zrt-, Irt-like protein domain-containing protein (ZIPCO) was suggested to import Fe^2+^ and Zn^2+^ into the cytosol and localized to the parasite plasma membrane (PPM) in *P. berghei* sporozoites in indirect immunofluorescence assays (34), but *Pf*ZIPCO (PF3D7_1022300) had not been studied yet. The chloroquine resistance transporter *Pf*CRT (PF3D7_0709000) and the natural resistance-associated macrophage protein *Pf*NRAMP (PF3D7_0523800, also called *Pf*DMT1 for divalent metal transporter 1, although this abbreviation is already in use for the drug/metabolite transporter 1) have been detected at the digestive vacuolar (DV) membrane (35, 36). Both proteins were proposed to export Fe^2+^ into the cytosol in symport with protons on the basis of transport assays using *Xenopus* oocytes (37) and proliferation assays with a conditional knockdown line under different iron conditions (38), respectively.

In *Saccharomyces cerevisiae*, a model organism for eukaryotic iron homeostasis, the mitochondrial carrier protein MRS3 (mitochondrial RNA-splicing protein 3) was shown to ensure Fe^2+^ supply to the mitochondrion (39–41) and its ortholog *Tg*MIT (mitochondrial iron transporter) was detected at the same organelle in *T. gondii*. (33). The mitochondrion of *P. falciparum* is also the focal point for cellular iron metabolism and contains iron-dependent proteins implicated in the biosynthesis of heme and iron-sulfur clusters, redox reactions, and electron transport (18). Because of sequence similarity (35.1% identity with the yeast ortholog, E = 3 x 10^-14^, 26% coverage), it was proposed that *Pf*MRS3 (also known as mitoferrin (*Pf*MFRN), PF3D7_0905200) mediates Fe^2+^ import into the mitochondrion in *P. falciparum* (42). However, no experimental evidence was collected and it is known that not only the localization but also the structure and function of homologous proteins can vary in related apicomplexan parasites (42, 43).

Despite the importance of iron for *P. falciparum* virulence, understanding of the molecular mechanisms of iron sensing, acquisition, utilization, and regulation in the parasite remains limited. The goal of this exploratory study was to dissect how the parasite responds to differences in iron availability in its environment and to identify putative iron transporters as potential new antimalarial drug targets. We investigated growth and gene expression of the laboratory *P. falciparum* strain 3D7 under control (iron-replete), high-iron and low-iron conditions, and in the presence of the iron-regulatory peptide hormone hepcidin. In the human body, hepcidin is produced to reduce the concentration of serum iron when it rises above a certain threshold. The hormone binds specifically to ferroportin on the surface of many cell types including erythrocytes (44) and can sterically inhibit the transporter’s iron export activity, thereby increasing intracellular and decreasing serum iron levels (45, 46). Here, whole-transcriptome sequencing was used to identify putative iron transport proteins on the basis of differential gene expression patterns between high vs. low-iron conditions. We then further studied these proteins by analyzing their subcellular localization in live parasites as well as their predicted 3D structures and by determining growth rates of the respective knockout or knockdown parasite lines under various conditions.

## RESULTS

### Elevated erythrocyte labile iron levels promote *P. falciparum* proliferation in vitro

To investigate whether labile iron levels in the erythrocyte correlate with parasite replication rates, we established different iron conditions in vitro. The first approach was to culture *P. falciparum* 3D7 parasites in 0 Rh+ erythrocytes from voluntary blood donations by Caucasians aged 18 to 21 at the University Medical Center Hamburg-Eppendorf in Germany. Therefore, samples from a person with an elevated ferritin level (greater than 200 µg/L (47), in this case 231 µg/L ferritin, 18.2 g/dL Hb, 51.5% hematocrit), an iron-deficient individual (serum ferritin < 12 ng/ml (48), here: 3 µg/L ferritin, 11.4 g/dL Hb, 36.3% hematocrit) and a healthy donor (21 µg/L ferritin, 15.0 g/dL Hb, 42.3% hematocrit) were used. Secondly, infection of red blood cells from other healthy individuals with or without the addition of 0.7 µM hepcidin to the culture medium was compared. This concentration was chosen as it had the strongest effect on parasite proliferation in preliminary experiments, and is expected to increase intracellular Fe^2+^ as it is twice as high as the hepcidin level needed to reduce ^55^Fe export from preloaded mature erythrocytes by 30% within one hour of incubation (44).

Relative labile iron levels in uninfected erythrocytes were estimated by determining the mean fluorescence intensity (MFI) of the iron-sensitive dye Phen Green SK in 100,000 cells per replicate using flow cytometry. As binding of ferrous iron to the metal-binding moiety causes fluorescence quenching of the fluorophore, a reduction in fluorescence intensity indicates higher labile iron levels (49). In erythrocytes from the iron-deficient donor, the Phen Green SK MFI was 43% higher relative to control, confirming reduced labile iron levels (Fig. 1A). The parasite replication rate after one intraerythrocytic developmental cycle (IDC) decreased by 16% (Fig. 1B), the DNA content of late schizonts by 19% (Fig. 1C) and the number of merozoites counted per late schizont by 14% (Fig. 1D). In contrast, labile iron levels of erythrocytes from the donor with higher iron status were only slightly increased (without statistical support, two-tailed unpaired *t* tests with Welch’s correction for unequal variances and adjusted with the Holm-Šídák method for multiple comparisons, *P* = 0.25) relative to blood with normal iron level (healthy control) – as were the parasite proliferation rate, the DNA content and the merozoite number of mature schizonts (Fig. 1). To further increase intracellular labile iron levels, we incubated parasites with 0.7 µM hepcidin during one IDC, resulting in 11% reduced Phen Green SK MFI compared to control (Fig. 1A). Under these conditions, the parasite growth rate increased by 57% (Fig. 1B), the DNA content per schizont by 16% (Fig. 1C), and the number of merozoites produced per schizont by 15% (Fig. 1D).

**Figure 1:**
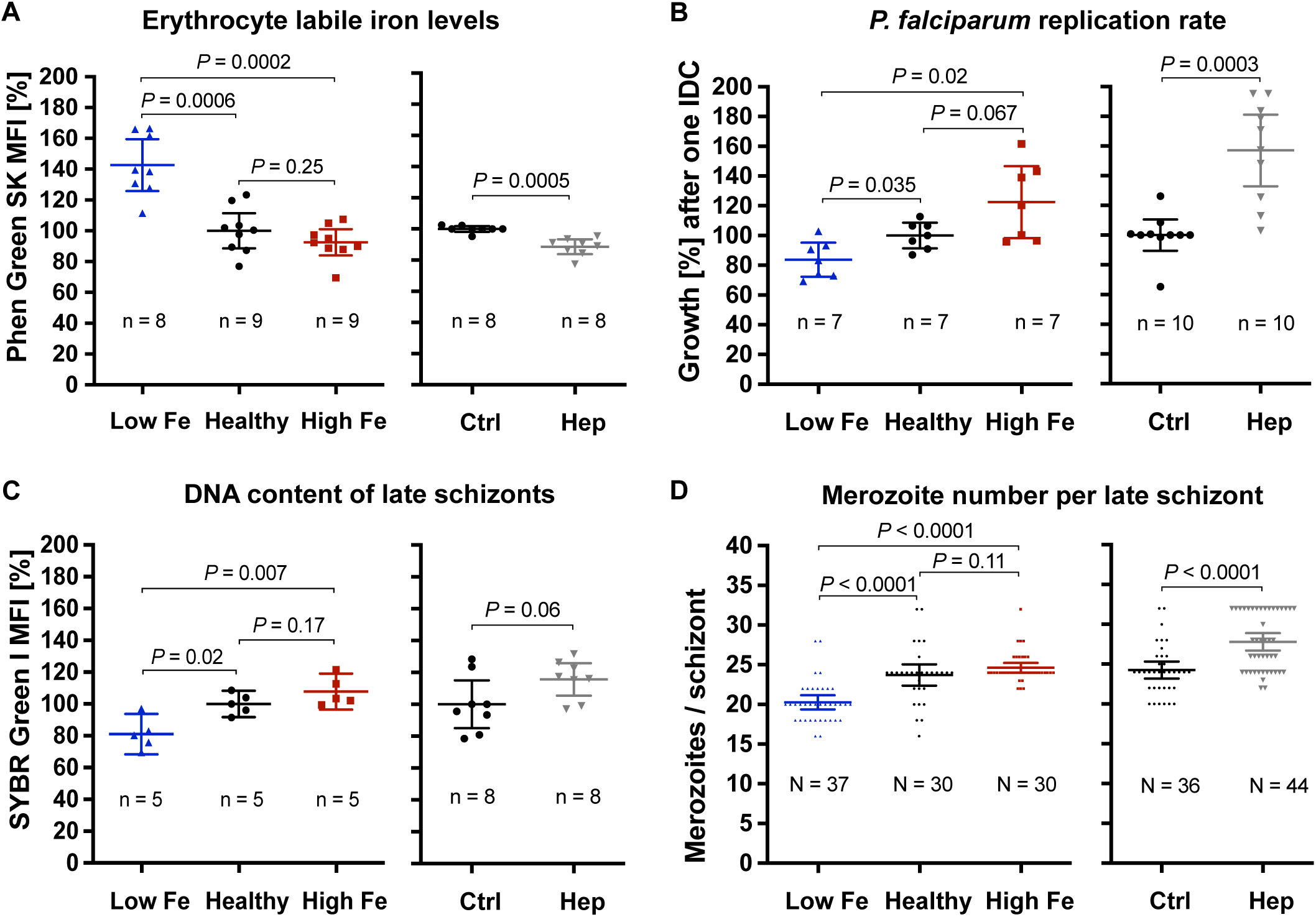
Effects of the iron status of the blood donor and of hepcidin on (A) erythrocyte labile iron levels, (B) *P. falciparum* 3D7 growth rates, (C) DNA content per mature schizont, and (D) the number of merozoites per mature schizont. The relative labile iron level and DNA content per cell were assessed in the presence or absence of 0.7 µM hepcidin (Hep) by measuring the mean fluorescence intensity (MFI) of Phen Green SK or SYBR Green I compared to control (Ctrl, untreated, normal hemoglobin level) using flow cytometry. Therefore, 100,000 cells were analyzed per replicate. Parasite growth rates refer to the fold change in parasitemia after one intraerythrocytic developmental cycle in vitro relative to control as determined by flow cytometry with SYBR Green I (104). Mature schizonts were obtained by treating schizonts at 40 hpi with 1 mM compound 2 (4-[7-[(dimethylamino)methyl]-2-(4-fluorphenyl)imidazo[1,2-α]pyridine-3-yl]pyrimidin-2-amine) for 8 h. To count the number of merozoites, Giemsa-stained blood smears were analyzed microscopically and only single-infected cells with one digestive vacuole were considered. Means and 95% confidence intervals (indicated by error bars) are shown. Statistical significance was calculated with two-tailed unpaired t tests with Welch’s correction for unequal variances and adjusted with the Holm-Šídák method for multiple comparisons except for merozoite numbers, which were compared using Mann-Whitney test. The number of independent experiments is represented as n and the number of parasites imaged in three independent experiments as N.

Taken together, these data show that parasites grown in erythrocytes from an iron-deficient donor displayed significantly reduced growth rates compared to healthy control. Our in vitro results also demonstrate that hepcidin treatment of control erythrocytes elevated intracellular Fe^2+^ concentrations and promoted parasite proliferation.

### RNA-sequencing reveals differential expression of putative iron transporters

To identify iron-regulated mechanisms and putative iron transporters in *P. falciparum*, we carried out whole-transcriptome profiling using bulk RNA-sequencing (Fig. 2). *P. falciparum* 3D7 parasites were cultured either using erythrocytes from a donor with higher, control (healthy) or low iron status (experiment 1); or with red blood cells from another healthy donor in the presence or absence of 0.7 µM hepcidin (experiment 2). Samples from three biological replicates per condition were harvested at the ring and trophozoite stage (6 – 9 and 26 – 29 hours post invasion, hpi) during the second IDC under the conditions specified.

**Figure 2:**
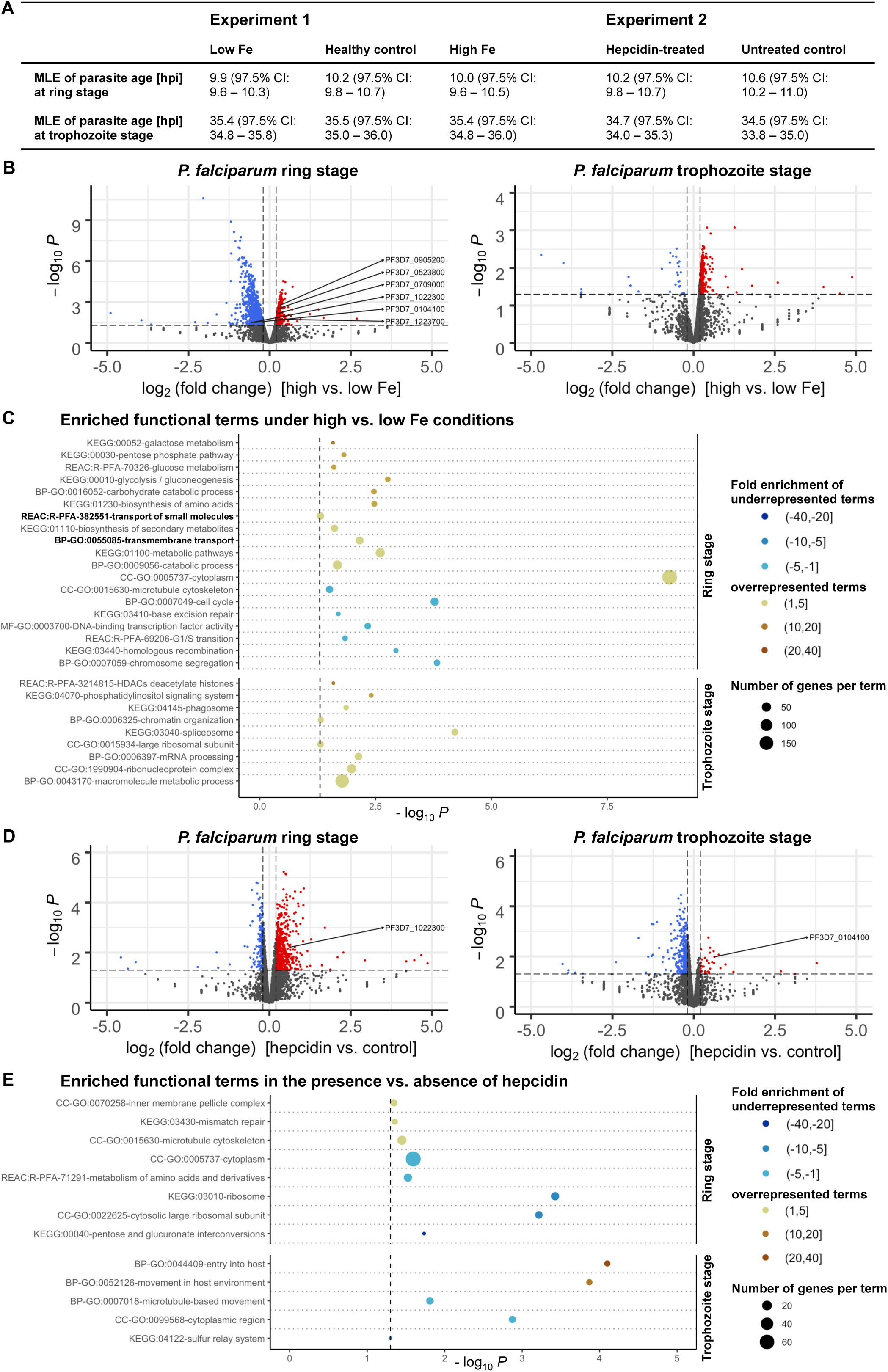
Differential expression of *P. falciparum* 3D7 genes under various iron conditions. Parasites were cultured with erythrocytes from an individual with high, medium (healthy) or low iron status (experiment 1) or with red blood cells from another healthy donor in the presence or absence of 0.7 µM hepcidin (experiment 2). Samples were harvested at the ring and trophozoite stage (6 – 9 and 26 – 29 hours post invasion, hpi) with three biological replicates per time point and condition. The maximum likelihood estimation (MLE) of the average developmental age of the parasites for each condition and time point (**A**) was calculated using an algorithm developed by Avi Feller and Jacob Lemieux (50). CI, confidence interval. The volcano plots (**B** and **D**) show transcriptional changes of all parasite genes. Red dots indicate significantly (*P* < 0.05, exact test for negative binomial distribution) upregulated genes (log_2_ (fold change) ≥ 0.2), blue dots stand for significantly downregulated genes (log_2_ (fold change) ≤ −0.2), while grey dots represent genes that did not significantly differ in transcription under the conditions described (*P* ≥ 0.05 and / or −0.2 < log_2_ (fold change) < 0.2). Differentially expressed genes encoding putative iron transporters (see Table 1) are labeled. Panels **C** and **E** show the enrichment of Gene Ontology (GO), Kyoto Encyclopedia of Genes and Genomes (KEGG) and Reactome (REAC) terms among significantly regulated genes excluding *var*, *stevor* and *rifin* gene families at the two time points. The functional terms were summarized using REVIGO (122) to remove redundancies, represented by circles and plotted according to the significance of their enrichment (-log_10_ (adjusted *P*), hypergeometric test). The size of the circle is proportional to the number of differentially regulated genes in the dataset that are associated with the respective term, while the color stands for the fold enrichment. The gray dashed line indicates the threshold of the adjusted *P* value (-log_10_ 0.05 = 1.3).

**Table 1:**
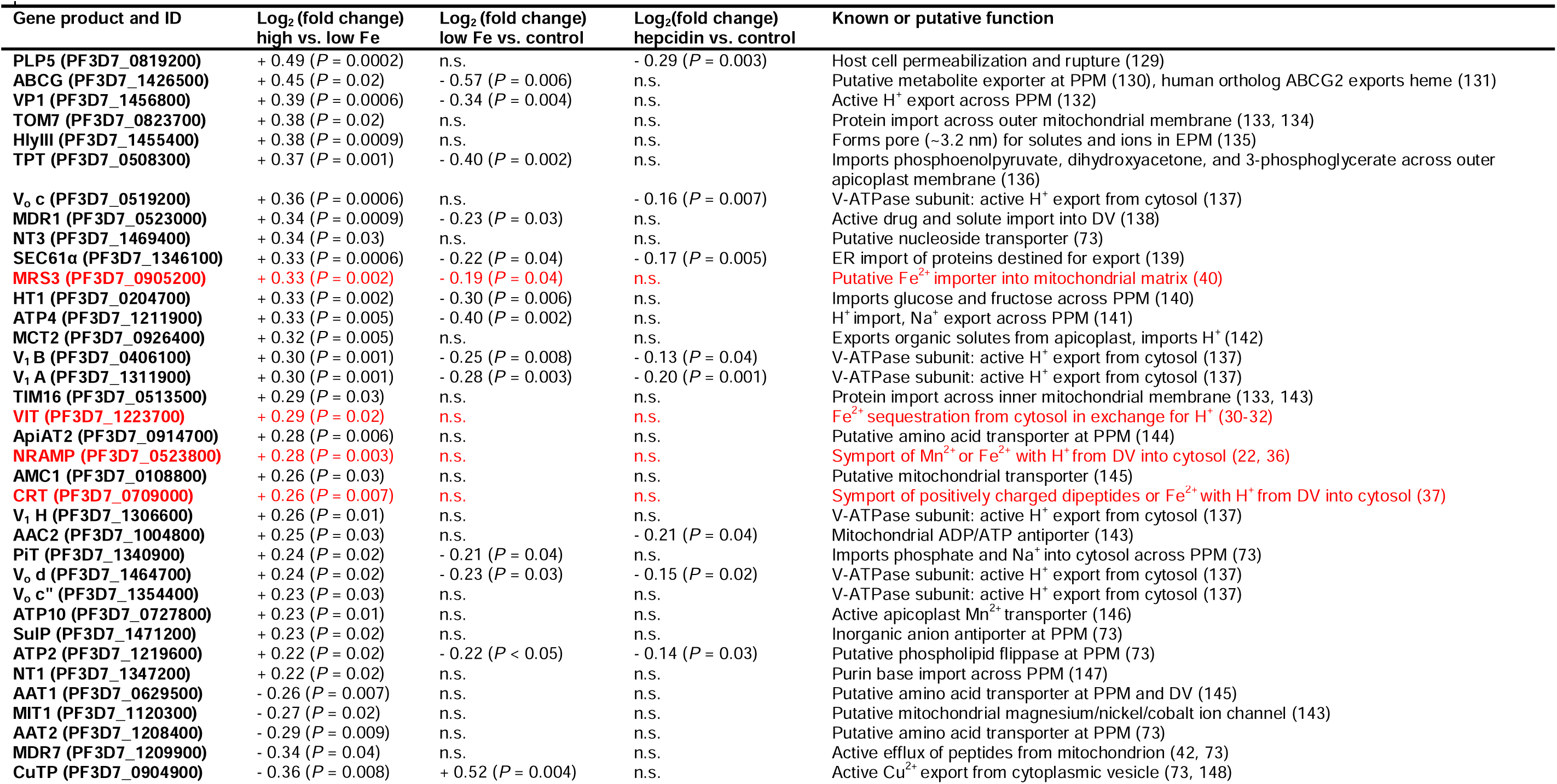
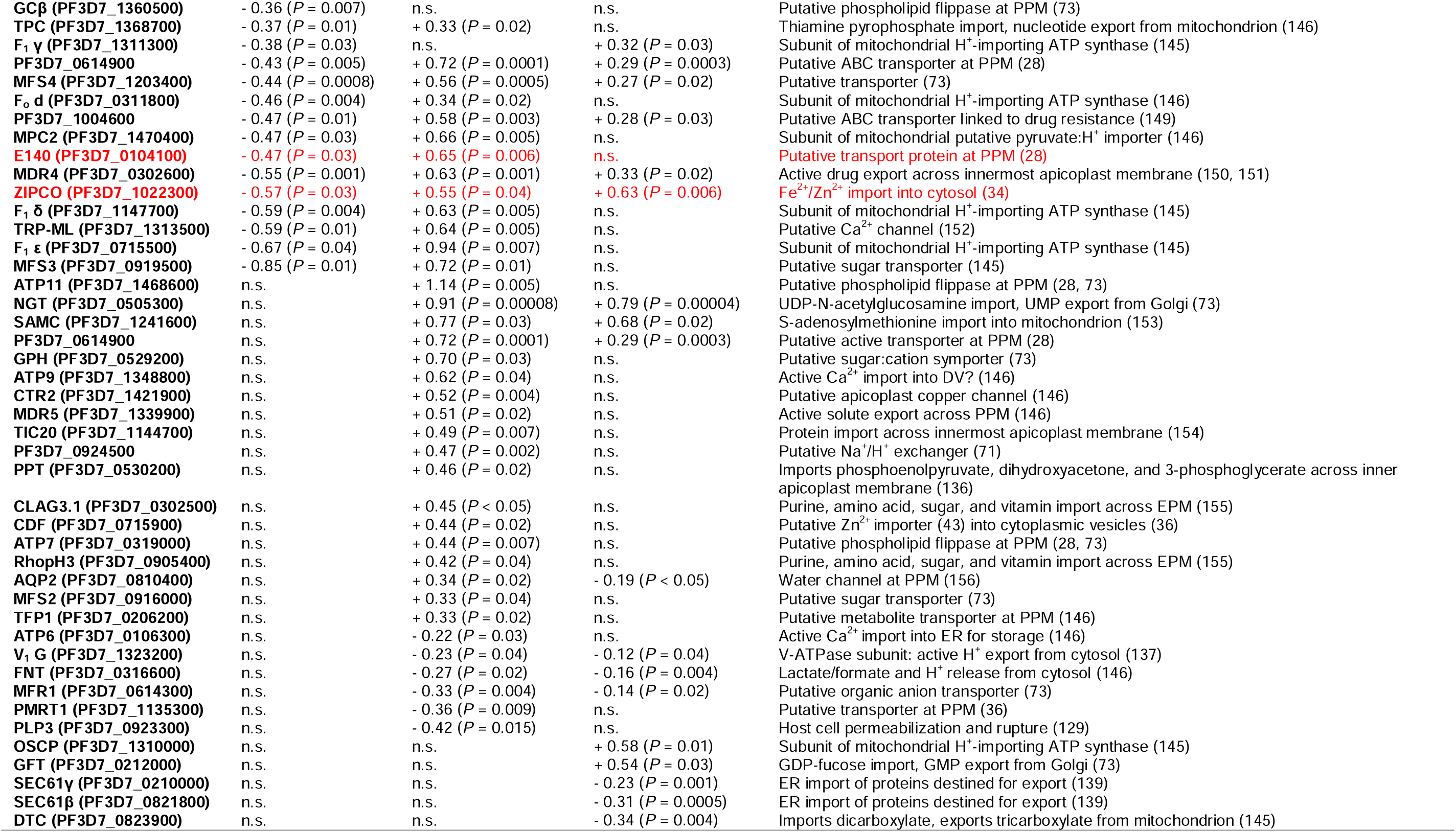
*P. falciparum* transport proteins with differential gene expression under various iron conditions. Putative and known transporter genes were filtered from differentially expressed genes in the described RNA-sequencing experiments using a list of *P. falciparum* transport proteins (28). The log_2_ (fold change) of gene expression at the ring stage (6 – 9 hours post invasion) and known or proposed functions are indicated for significantly regulated genes (exact *P* < 0.05). The identified (potential) iron transport proteins are highlighted in red. DV, digestive vacuole; EPM, erythrocyte plasma membrane; PPM, parasite plasma membrane.

To exclude the possibility that differences in mRNA abundance were a consequence of divergent progression through the IDC under different nutritional conditions, we assessed the average developmental age of the parasites in each sample on the basis of a statistical maximum likelihood estimation (MLE) method of transcriptional patterns according to Lemieux et al. (50). The general transcriptional patterns of parasites were highly similar at individual time points and consistent across different experimental treatments, corresponding to those of a 3D7 reference strain (51) at approximately 10 hpi and 35 hpi (Fig. 2A). This indicates that differences in mRNA abundance of parasites were not caused by divergent progression through the IDC but by direct effects of the experimental treatments. As the 3D7 strain we used for the experiments had a reduced total IDC length of 44 h instead of 48 h, possibly because of gene deletions that may have occurred during long-term culturing (52, 53), it progresses through the cycle faster than the 3D7 reference strain (51). This may explain why the calculated MLEs of parasite age were higher than the actual values of 6 – 9 hpi and 26 – 29 hpi (Fig. 2A).

Using a threshold of 1.5 for the fold change (FC) in gene expression (log_2_ FC of 0.585 or −0.585) yielded twelve significantly upregulated and 175 downregulated genes in ring-stage parasites under high vs. low-iron conditions (*P* < 0.05, exact test for the negative binomial distribution with Benjamini-Hochberg correction (54)). As differences in transporter gene transcription are typically small (55, 56), we examined the 351 upregulated and 770 downregulated genes with a significant expression change and a minimum absolute value of the log_2_ FC of 0.2 for this comparison (Fig. 2B). The full RNA-sequencing datasets are available in the BioStudies repository (57) under accession number E-MTAB-13411 (https://www.ebi.ac.uk/biostudies/studies/E-MTAB-13411) and differential gene expression test results for individual genes are shown in Supplementary Tables S1 and S2. The highly polymorphic *var*, *stevor*, and *rifin* gene families were excluded from downstream analyses because of their significant sequence diversity between parasites of the same strain during mitotic growth (58, 59). Functional Gene Ontology (GO), Kyoto Encyclopedia of Genes and Genomes (KEGG) and Reactome (REAC) term enrichment analyses of differentially expressed genes (DEGs) were performed using the g:Profiler web server (60).

Under high vs. low-iron conditions at the ring stage (6 – 9 hpi), the GO term for biological process GO:0055085 “transmembrane transport” was 2.8-fold enriched (*P* = 0.007, hypergeometric test) among significantly upregulated parasite genes (Fig. 2C). Using the recently updated *P. falciparum* transporter list (28), all genes with differential expression levels at the ring stage were then screened for transport proteins and all of the five genes previously proposed as iron transporters in *Plasmodium* (VIT, ZIPCO, NRAMP/DMT1, CRT, MRS3/MFRN (18, 43)) were found differentially expressed (Table 1). Other significantly enriched functional terms at the ring stage under high-iron conditions were GO:0009056 “catabolic process”, GO:0020020 “food vacuole”, KEGG:01100 “metabolic pathways”, and GO:0005737 “cytoplasm” (Fig. 2C). Among downregulated genes under high vs. low-iron conditions at the ring stage, KEGG:03440 “homologous recombination”, KEGG:03410 “base excision repair”, GO:0007049 “cell cycle”, and GO:0015630 “microtubule cytoskeleton” were enriched (Fig. 2C). At the more metabolically active trophozoite stage (26 – 29 hpi), processes related to mRNA splicing and protein production were overrepresented in upregulated genes, as indicated by the 2.9-fold enrichment (*P* = 0.00006) of the KEGG:03040 pathway “spliceosome” and the 2.5-fold enrichment (*P* < 0.05) of the GO:0015934 term “large ribosomal subunit” (Fig. 2C).

In contrast, hepcidin treatment resulted in reduced metabolism compared to control conditions, as KEGG:00040 “pentose and glucuronate interconversions”, REAC:R-PFA-71291 “metabolism of amino acids and derivatives”, GO:0005737 “cytoplasm”, and GO:0015934 “large ribosomal subunit” were significantly enriched in downregulated genes during the parasite ring stage at 6 – 9 hpi (Fig. 2E). Among significantly upregulated genes in the presence vs. absence of hepcidin, the terms GO:0070258 “inner membrane pellicle complex” (*P* < 0.05), KEGG:03430 “mismatch repair” (*P* = 0.04), and GO:0015630 “microtubule cytoskeleton” (*P* = 0.04) were enriched at the ring stage. GO:0044409 “entry into host” (*P* = 0.00008) and GO:0052126 “movement in host environment” (*P* = 0.0001) were overrepresented at the trophozoite stage (Fig. 2E), possibly linked to the observed increase in parasite proliferation (Fig. 1B).

Our RNA-sequencing data also revealed the differential expression of genes involved in epigenetic, transcriptional, translational, and post-translational regulation. Under high vs. low-iron conditions, histone deacetylation and chromatin organization processes as well as GO:1990904 “ribonucleoprotein complex” were significantly enriched in upregulated genes at the trophozoite stage, and GO:000370 “DNA binding transcription factor activity” in downregulated genes at the ring stage (Fig. 2C). Furthermore, the known iron-regulatory protein *Pf*IRP or aconitate hydratase (61, 62) was upregulated during the ring stage under high vs. low-iron conditions (log_2_ FC = 0.49, *P* = 0.00003) and downregulated in the presence of hepcidin (log_2_ FC = −0.27, *P* = 0.01) as compared to control. Many protein kinases involved in post-translational modifications and endocytosis were also upregulated at 26 – 29 hpi at high vs. low iron levels, as indicated by the enriched terms GO:0043170 “macromolecule metabolic process” and KEGG:04070 “phosphatidylinositol signaling system” (Fig. 2C).

### Localization of putative iron transporters in *P. falciparum*

On the basis of transcriptomic profiles and the *P. falciparum* transporter list (28), six proteins with a potential role in iron transport were identified (Table 1 and 2). The subcellular localization of the four proteins that had not yet been localized in *P. falciparum* (*Pf*MRS3, *Pf*VIT, *Pf*ZIPCO, and *Pf*E140) was then examined by endogenous tagging with GFP and confocal imaging of live parasites under physiological control conditions. At least two cell lines were generated per transporter candidate with consistent results and representative example images are shown in Fig. 3. Diagnostic PCRs confirmed the fusion of *gfp* to the respective gene of interest and the absence of parental DNA at the original locus (Supplementary Fig. S2). Only the *Pf*MRS3 reporter cell line still contained wild-type DNA of the parental parasites even after prolonged WR99210/neomycin selection and limiting dilution cloning (Supplementary Fig. S2), indicating the importance of this mitochondrial transporter for asexual parasite growth during the blood stage.

**Figure 3:**
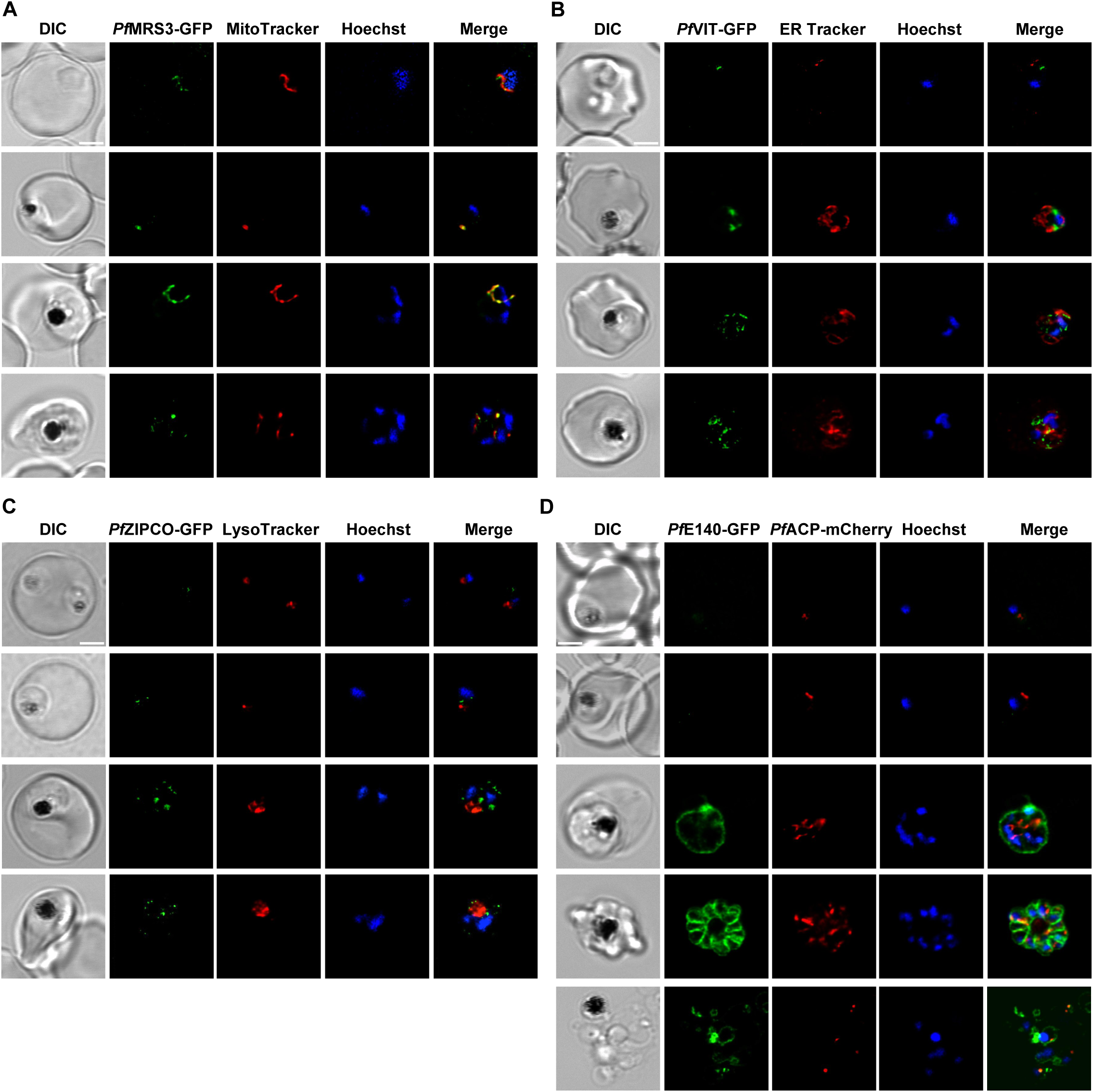
Subcellular localization of known and putative iron transport proteins. Representative erythrocytes infected with *P. falciparum* 3D7 parasites endogenously expressing GFP-tagged *Pf*MRS3 (**A**), *Pf*VIT (**B**), *Pf*ZIPCO (**C**) or *Pf*E140 (**D**) were additionally stained with the fluorescent dyes Hoechst-33342, MitoTracker Red, ER Tracker Red and/or LysoTracker Deep Red. Co-transfection with a construct that encodes the 60 N-terminal amino acids of acyl carrier protein (*Pf*ACP) tagged with mCherry (125) resulted in red fluorescence of the apicoplast. Live-cell images were taken under physiological conditions at 37°C using an SP8 confocal laser-scanning microscope (Leica). DIC, differential interference contrast. Scale bar, 2 µm.

The GFP-tagged mitochondrial carrier protein *Pf*MRS3 exclusively localized to the mitochondrion, as determined by colocalization with MitoTracker Red (Fig. 3A, Supplementary Video S1). *Pf*VIT-GFP displayed a punctate fluorescence pattern within the cytoplasm (Fig. 3B, Supplementary Video S2), resembling that of *Pf*ZIPCO-GFP (ZIP domain-containing protein, PF3D7_1022300, Fig. 3C, Supplementary Video S3). These structures did not colocalize with ER Tracker Red in live cells (Fig. 3B, Supplementary Video S2). For both *Pf*VIT-GFP and *Pf*ZIPCO-GFP, the number of cytoplasmic foci increased as the parasites matured from the ring to the late schizont stage (Fig. 3B and C). To test whether these could be acidocalcisomes, we employed LysoTracker Deep Red, commonly used to visualize small acidic organelles in *T. brucei* (24). However, the fluorescent dye only stained the DV in *P. falciparum* (Fig. 3C, Supplementary Video S3) and no acidocalcisome-specific marker is currently available for this parasite.

GFP-tagged *Pf*E140 (PF3D7_0104100), also known as conserved *Plasmodium* membrane protein or CPMP (63), localized to the parasite plasma membrane, as evidenced by the ring-like fluorescence pattern around newly formed merozoites (Fig. 3D, Supplementary Video S4). The fluorescence intensity was very low at the ring and early trophozoite stage compared to schizonts. Because of amino acid sequence similarity (E = 9 x 10^-5^, 22.5% identity, 66% coverage) to the essential apicoplast transporter *Pf*DER1-2 (29, 64), we also investigated the potential colocalization with the apicoplast marker *Pf*ACP (acyl carrier protein), which could not be detected (Fig. 3D, Supplementary Video S4).

### The role of *Pf*VIT, *Pf*ZIPCO and *Pf*E140 for asexual parasite growth

To study the function of the putative transport proteins identified, we used targeted gene disruption (TGD) by selection-linked integration (SLI) to generate the corresponding knockout parasite lines for the putative iron transporters that are non-essential during *P. falciparum* blood stage: *Pf*VIT and *Pf*ZIPCO (Fig. 4A, Supplementary Fig. S2). As GFP was cloned in frame with the truncated version of the respective transporter (the N-terminal 143 amino acids (aa) of 274-aa *Pf*VIT or 117 of the 325 aa of *Pf*ZIPCO), the subcellular localization of the resulting GFP fusion protein was also assessed. *Pf*VIT(1–143)-GFP localized to cytoplasmic structures and *Pf*ZIPCO(1–117)-GFP to the DV and cytoplasmic vesicles (Fig. 4A). Proliferation assays were then performed to determine the importance of the respective transporter for parasite growth. While the *Pf*VIT knockout had no effect on parasite growth under standard conditions, addition of hepcidin reduced the growth rate of the ΔVIT line by 30% (Fig. 4B). Of note, hepcidin generally had a smaller effect after two cycles of incubation (Fig. 4B) than after one cycle compared to the first IDC (Fig. 1B). Unexpectedly, knocking out *Pf*ZIPCO led to a growth rate increase by 42% after two IDCs relative to wild-type 3D7 parasites (Fig. 4B).

**Figure 4:**
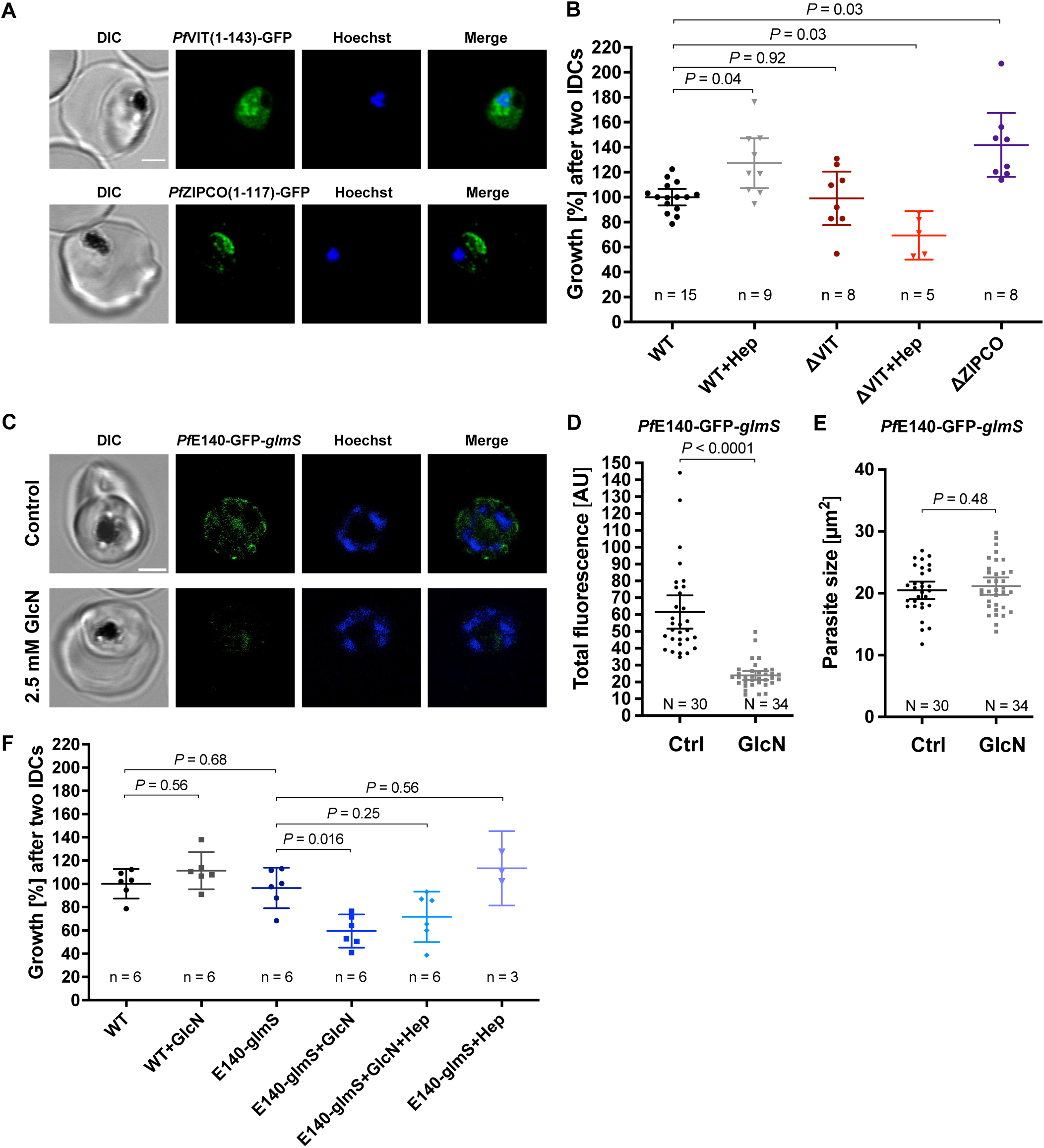
*Pf*VIT and *Pf*E140 are important for *P. falciparum* growth and may be involved in intracellular iron homeostasis. **A** Representative erythrocytes infected with *P. falciparum* 3D7 parasites that endogenously express a truncated version of *Pf*VIT or *Pf*ZIPCO tagged with GFP (green). **B** Growth rates of knockout parasite lines generated. **C** Reduction of *Pf*E140-GFP fluorescence (green) in live 3D7 parasites caused by *glmS*-mediated knockdown that was induced by treatment with 2.5 mM glucosamine (GlcN) for 36 h compared to untreated control (Ctrl). **D** Total parasite fluorescence intensities were quantified as background-corrected integrated densities using ImageJ version 2.9.0/1.53t (110) and compared using Mann-Whitney test. **E** The size of the parasites was measured as the area of the region of interest and compared using equal variance unpaired t test. **F** Conditional knockdown of *Pf*E140 induced by treatment with 2.5 mM GlcN results in a growth defect during asexual blood stage development. Live parasites were stained with Hoechst-33342 (blue) and imaged under physiological conditions at 37°C using an SP8 confocal laser-scanning microscope (Leica). DIC, differential interference contrast. Scale bar, 2 µm. Error bars represent 95% confidence intervals of the mean, N the number of parasites analyzed, n the number of independent experiments and Hep treatment with 0.7 µM hepcidin. Growth rates refer to the fold change in parasitemia after two intraerythrocytic developmental cycles in vitro relative to untreated wild-type 3D7 parasites (WT) as determined by flow cytometry with SYBR Green I (104). Statistical significance of growth differences was calculated with two-tailed unpaired t tests with Welch’s correction for unequal variances and adjusted with the Holm-Šídák method for multiple comparisons.

*Pf*E140 is predicted to be essential (65) and the only putative iron transporter identified that localized to the PPM (Fig. 3), thus potentially important for iron uptake in *P. falciparum*. For an inducible knockdown, a *glmS* ribozyme sequence (66) was introduced upstream of the 3’ untranslated region in the pSLI plasmid, allowing for conditional mRNA degradation by adding 2.5 mM glucosamine (GlcN) to the culture medium (Fig. 4C, Supplementary Fig. S2). The knockdown led to a 61% decrease in total parasite fluorescence intensity after 36 hours of GlcN treatment (Fig. 4D) without affecting parasite size compared to untreated control (Fig. 4E). Addition of GlcN also caused a 38% growth rate reduction of the *Pf*E140-GFP-*glmS* line, which was rescued by hepcidin treatment to a proliferation level that was not significant different from that under standard culture conditions (*P* = 0.25, Fig. 4F). The generation of a *Pf*MRS3-knockdown line was not successful after four independent attempts, supporting the essentiality of the gene for asexual growth (65).

### Characterization and functional implications of predicted protein structures

We next took advantage of the recent progress in protein structure prediction and generated models of the putative iron transport proteins identified (Table 2) using AlphaFold2 (67, 68) and AlphaFold2-multimer (69). The transmembrane regions of the proteins typically exhibited the highest confidence score, while some other protein portions appeared unstructured (Fig. 5A). Regions that are likely located within a membrane were validated by inspecting the molecular lipophilicity potential of the protein surfaces (Fig. 5B). A clear hydrophobic belt was observed for all proteins and their orientation in the membrane was determined on the basis of orthologous proteins. As transport cavities with negatively charged residues are a hallmark of heavy metal ion transporters, we analyzed the distribution of charge on the surface of the proteins and looked for negatively charged regions to assess the capacity to bind cations like Fe^2+^ (Fig. 5C). To gain further insights into the functions of the proteins identified, we also compared the predicted structures with those of well-characterized homologs from *S. cerevisiae*, *Eucalyptus grandis, Bordetella bronchiseptica* and *Staphylococcus capitis* (Table 2, Supplementary Fig. S3 and S4).

**Figure 5:**
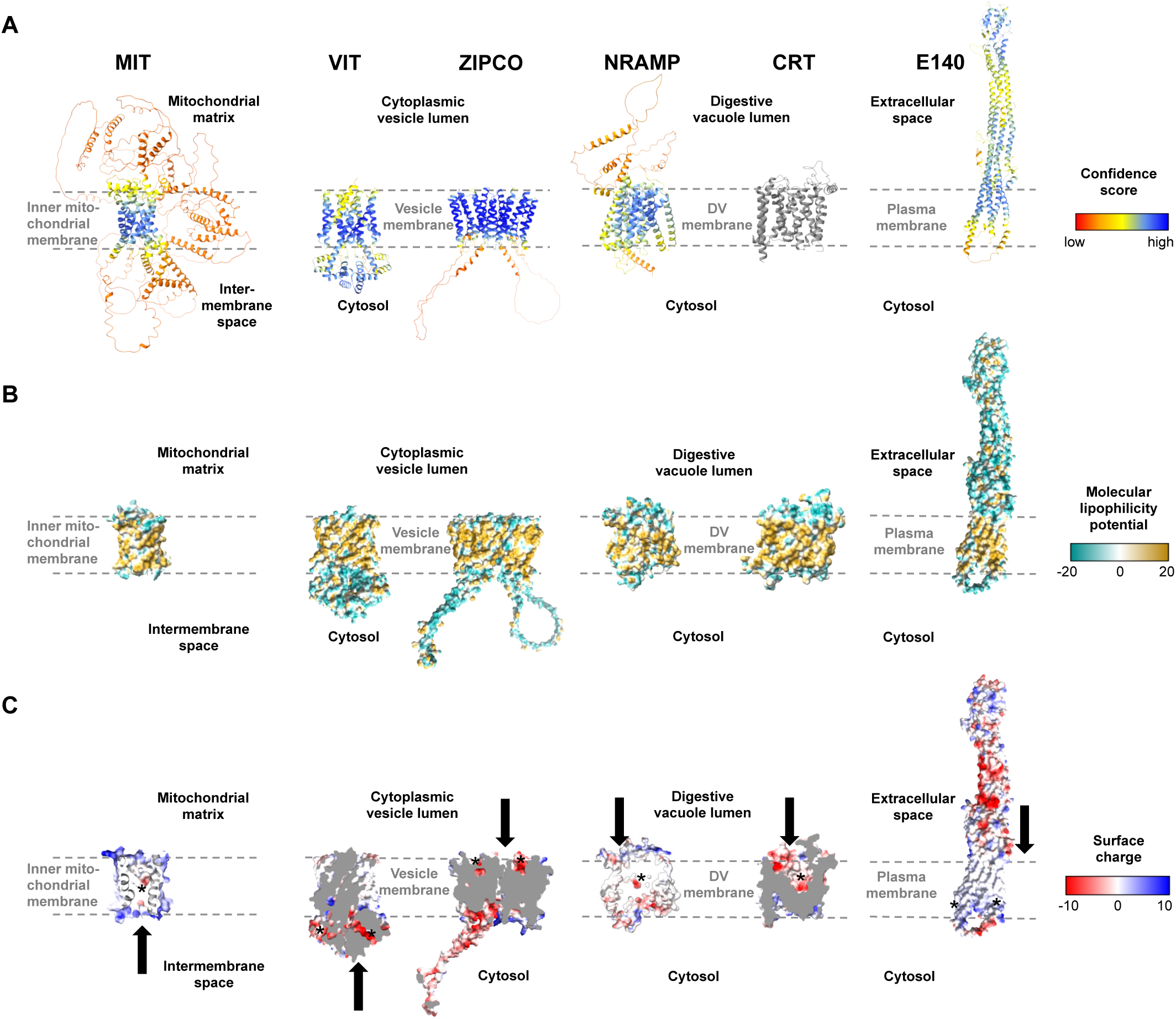
Structures of known and putative *P. falciparum* iron transporters as viewed from the membrane plane. **A** Predicted protein structures with per-residue pLDDT (predicted local distance difference test) confidence scores on a scale from 0 to 100, where blue represents high and red low confidence, respectively. The experimentally determined structure of *Pf*CRT is shown in gray. **B** Molecular lipophilicity potential of the protein surfaces as implemented in UCSF ChimeraX; tan is hydrophobic and cyan hydrophilic. Dashed lines above and below the tan regions of all proteins indicate the respective membrane and disordered loops were removed for clarity. **C** Surface charge of the proteins with positively charged areas colored blue and negatively charged ones red. Putative cation-binding site are indicated with an asterisk and transport directions by arrows. *Pf*E140 likely forms a dimer but is shown as a monomer, as no predicted dimer structure could be obtained using AlphaFold2-multimer. The putative cation-binding sites for this protein are based on DeepFRI gradCAM scores for the functional term GO:0015075 “monoatomic ion transmembrane transporter activity” (Supplementary Fig. S5).

**Table 2:**
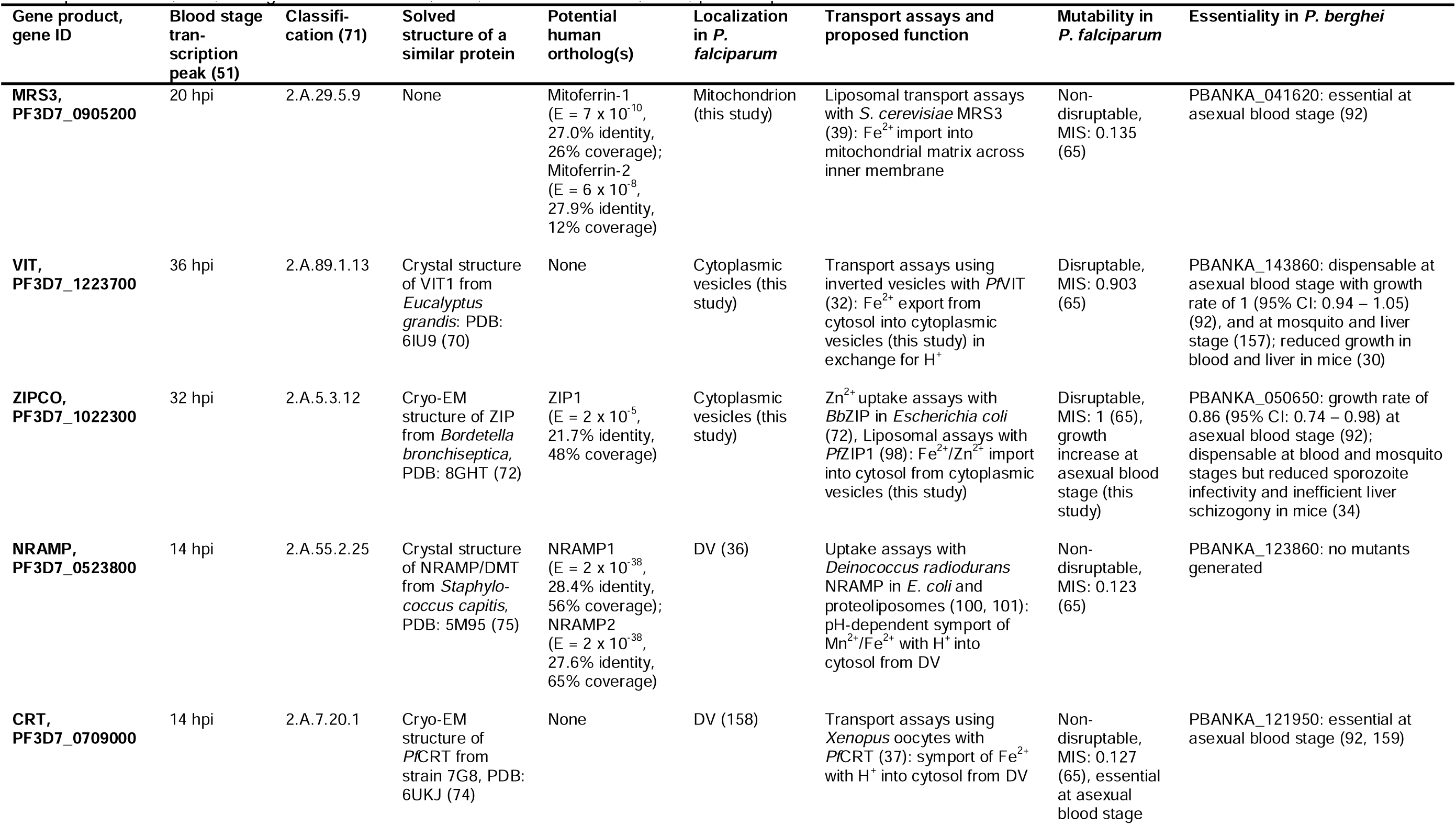
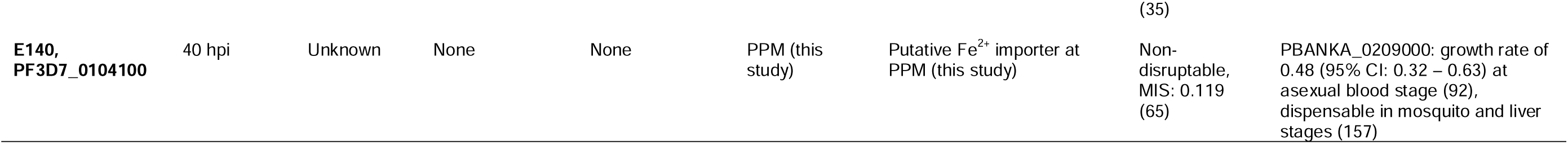
Proteins identified by RNA-sequencing that may be involved in iron transport in *P. falciparum.* The classification of the proteins identified is indicated according to the Transport Classification Database (71). Data on human orthologs was retrieved using the NCBI position-specific iterated (PSI) BLAST with default settings at https://blast.ncbi.nlm.nih.gov/Blast.cgi (29). DV, digestive vacuole; E, expect value; EM, electron microscopy; hpi, hours post invasion; MIS, mutagenesis index score; PDB, Protein Data Bank; PPM, parasite plasma membrane.

The outer surface of *Pf*MRS3 (transport classification (TC): 2.A.29, mitochondrial carrier family) is positively charged (Fig. 5C) and there is a clear negatively charged patch in the putative binding pocket facing the mitochondrial intermembrane space. We compared the predicted *Pf*MRS3 structure with that of *S. cerevisiae* MRS3, which is known to import ferrous iron into the mitochondrial matrix across the inner membrane (39–41). The predicted structures of *Pf*MRS3 and *S. cerevisiae* MRS3 were superimposed with an average root mean square deviation of Cα atoms (Cα RMSD) of the 205 matched residues of 2.3 Å (Supplementary Fig. S3A and S4A). The conserved histidine residues His^48^ and His^105^ that were required for Fe^2+^ transport by *S. cerevisiae* MRS3 in reconstituted liposomes (39) are also present in *Pf*MRS3 and the three functionally relevant histidine residues identified in yeast are in a similar functional context in both structures (Supplementary Fig. S3A and S4A). This suggests that MRS3 may elicit similar molecular functions in *S. cerevisiae* and *P. falciparum*.

*Pf*VIT is highly similar to VIT1 from *E. grandis* (E = 7 x 10^-27^, 30.3% identity, 84% coverage), for which an experimental structure is available (Protein Data Bank (PDB) identifier: 6IU4). The plant protein crystallized as a homodimer (70), and the same oligomeric state was suggested for the vacuolar iron transporter family (TC: 2.A.89) protein in *P. falciparum* (31). A *Pf*VIT monomer also has five transmembrane domains and comprises a negatively charged region facing the cytosol that may enable cation transport (Fig. 5C). In agreement with this, one Fe^2+^ ion and two Zn^2+^ ions were bound by a strongly charged region on the cytosolic side of the *E. grandis* VIT1 monomer (70) and a highly similar putative binding pocket is present in the parasite protein (Supplementary Fig. S4B). In the structural alignment, 219 of the 227 residues of the experimental *Eg*VIT1^23-249^ structure are within 5 Å of the predicted structure of *Pf*VIT with an average Cα RMSD of 1.9 Å and the key residues in the metal-binding domain (Glu^102^, Glu^105^, Glu^113^, Glu^116^, using *Eg*VIT1^23-249^ numbering) are placed in a similar molecular context in the predicted structure of *Pf*VIT (Supplementary Fig. S3B and S4B). The residues in the transmembrane domain that are in the vicinity of the Co^2+^ ion in the *Eg*VIT1^23-249^ structure (Met^80^ and Asp^43^) are also conserved (Supplementary Fig. S3B), which is in line with a similar function of *Pf*VIT and *Eg*VIT1.

*Pf*ZIPCO contains seven transmembrane domains and was modeled as a homodimer (Fig. 5A), as it is part of the zinc (Zn^2+^)-iron (Fe^2+^) permease (ZIP) family (TC: 2.A.5), whose members usually function as homo- or heterodimers (71). The negatively charged patch in each binding pocket facing the vesicle lumen (Fig. 5C) may be involved in cation transport to the cytosolic side. In an overlay of the *Pf*ZIPCO model with the cryo-EM structure (PDB: 8GHT) of a ZIP transporter from *B. bronchiseptica* in the presence of either Zn^2+^ or Cd^2+^ ions (72), the average Cα RMSD of 140 sequence-aligned residues was 2.0 Å (Supplementary Fig. S3C and S4C). Several key residues of the metal binding site M1 of *Bb*ZIP (Met^99^, His^177^, Glu^181^, Glu^211^) were also found in *Pf*ZIPCO, whereas others (Asn^178^, Gln^207^, Asp^208^, Glu^240^) were different (Supplementary Fig. S3C), possibly resulting in divergent substrate specificity.

*Pf*NRAMP (TC: 2.A.55, metal ion (Mn^2+^-iron) transporter family) is a homolog of the human endosomal Fe^2+^ transporter 2/DMT1 (73) and contains twelve transmembrane domains (Fig. 5A). Like *Pf*CRT (TC 2.A.7, drug/metabolite exporter family), for which a recent cryo-EM structure (PDB: 6UKJ) is available (74), the predicted structure possesses a negatively charged region within its binding pocket facing the digestive vacuolar lumen (Fig. 5C). This is consistent with binding of cations such as Fe^2+^. The *Pf*NRAMP model was superimposed on the solved crystal structure of *S. capitis* NRAMP/DMT (PDB: 5M95, E = 1 x 10^-32^, 26.3% identity, 60% coverage), which was shown to bind Mn^2+^, Fe^2+^, Co^2+^, Ni^2+^, Cd^2+^ and Pb^2+^ (75). In the overlay, the average Cα RMSD of the 349 matched residues was 1.6 Å and the negatively charged cavity of *Pf*NRAMP was in close proximity to the Mn^2+^ ion bound to *S. capitis* NRAMP (Supplementary Fig. S3D and Supplementary Fig. S4D). Two of the four key residues required for ion coordination in the binding pocket of the bacterial protein (Asn^52^ and Asp^49^) are present in *Pf*NRAMP, whereas the two other residues (Met^226^ and Ala^223^) are changed to serine (75). While potential effects of these differences on substrate specificity and / or transport activity remain to be elucidated, *Pf*NRAMP is likely to perform cation transport from the DV into the cytosol.

*Pf*E140 is predicted to be anchored in the parasite plasma membrane by a bundle of five transmembrane domains (76) and forms a coiled coil with a hydrophilic region that displays negatively charged patches exposed to the extracellular side (Fig. 5A and B). No human orthologs could be identified for this highly conserved *Plasmodium* protein (29). As there is no obvious channel or cavity in the transmembrane region of the *Pf*E140 monomer (Fig. 5B and C), the helical bundles may form a dimer to enable ion transport. However, we were not able to obtain a *Pf*E140 dimer model with AlphaFold2-multimer because of its sequence length. To predict functional residues on the basis of the amino acid sequence and the AlphaFold2 structure of *Pf*E140, we used DeepFRI graph convolutional network (77), which has significant denoising capability and can reliably assign GO terms to residues in the protein. In particular, the terms GO:0022857 “transmembrane transporter activity” (DeepFRI gradCAM score 0.94), GO:0015075 “monoatomic ion transmembrane transporter activity” (score 0.78), and GO:0046873 “metal ion transmembrane transporter activity” (score 0.67) were assigned to a putative transmembrane region of *Pf*E140 with high confidence (Supplementary Fig. S5). We thus speculate that the protein is a transporter of metal ions.

## DISCUSSION

Here, we studied the role of iron in growth and transcription of *P. falciparum* by using blood from individuals of different iron status and by adding hepcidin as an iron-regulatory hormone and ferroportin inhibitor. Overall, our data demonstrate the importance of Fe^2+^ in parasite replication and development and highlight areas for further study. We showed that in vitro growth rates of *P. falciparum* 3D7 and the number of merozoites formed per schizont were reduced within erythrocytes that contain lower concentrations of labile iron, while culturing in blood from an individual with higher iron status did not lead to a significant increase in labile iron levels within erythrocytes or in parasite growth relative to control (Fig. 1). Consistent with this, reduced propagation of *P. falciparum* 3D7, Dd2, and FCR3-FMG was reported when erythrocyte samples from iron-deficient individuals used for parasite culture (10, 78). This effect was eliminated after these donors were iron-supplemented, whereas supplementation of healthy (iron-replete) donors did not significantly promote parasite growth (10). The strong increase in parasite replication in the presence of hepcidin relative to control conditions (Fig. 1B) may have been a result of enhanced invasion efficiency in addition to the increased number of merozoites formed (Fig. 1D). Earlier studies also found that higher hepcidin levels in blood samples were associated with elevated *P. falciparum* growth rates in vitro (78) and severe malaria in vivo (79), however, the effect of experimental hepcidin addition on parasite growth had not been assessed previously.

To identify putative iron transporters and iron-regulated processes, we carried out RNA-sequencing analyses of *P. falciparum* during the ring (6 – 9 hpi) and trophozoite (26 – 29 hpi) stages cultured under the different iron conditions described above. A higher number of biological processes and pathways were significantly enriched among DEGs when erythrocytes from donors with different iron status were used for parasite culture (Fig. 2C) compared to red blood cells from the same healthy donor in the presence vs. absence of hepcidin (total of 28 vs. 13 functional terms, Fig. 2E). This may reflect greater differences in the culture conditions; for instance, blood from the donor with high serum ferritin and Hb levels may have also contained more glucose or copper (80, 81), potentially explaining the more diverse physiological response of the parasite. Including erythrocyte samples from more individuals in the growth experiments and RNA-sequencing analysis would have provided further insights, however, the provision of sufficient blood from iron-deficient donors is limited by ethical constraints.

The availability of additional nutrients likely resulted in increased endocytosis and digestion of host cell contents in the DV of the parasite, leading to enhanced metabolism, mRNA splicing, and protein production. Interestingly, the terms KEGG:01100 “metabolic pathways” and GO:0005737 “cytoplasm” were also found to be enriched in upregulated parasite genes in children with high vs. low parasitemia (82, 83). RNA binding and mRNA splicing processes were previously reported to be overrepresented in upregulated genes in severe malaria linked to high parasite density (82–84). Hence, an increase in overall parasite fitness under high vs. low-iron conditions may explain the increase in parasite multiplication (Fig. 1B) and could be associated with higher parasitemia and disease severity. Consistent with the observed upregulation of transmembrane transporters at the parasite ring stage (6 – 9 hpi) under high vs. low-iron conditions, Mancio-Silva et al. found that the functional term “ion transporter activity” was enriched in *P. berghei* genes that were downregulated under caloric restriction at 6 and 10 hpi (16). Thus, transmembrane transporter genes may need to be transcribed at the beginning of the IDC to ensure that the appropriate level of transport proteins is available for nutrient acquisition and metabolite efflux during the subsequent metabolically active trophozoite and schizont stages.

Hepcidin plays a central role in mammalian iron homeostasis and reduces serum iron concentrations (85). It is also known that hepcidin levels are elevated in *P. falciparum*-infected individuals, especially those with high parasitemia (79, 86), and that malaria causes iron deficiency (79). The transcription profile of parasites treated with 0.7 µM hepcidin showed similarities to those cultured in erythrocytes from the iron-deficient donor compared to standard conditions in terms of downregulated catabolic and translation processes as well as transport protein regulation (Fig. 2, Table 1). This may be related to the fact that an aberrant hepcidin increase causes systemic iron deficiency as a result of restricted iron availability (87). The upregulation of genes involved in merozoite motility (*Pf*MTIP, *Pf*GAP45, and various inner membrane complex proteins) and host cell entry (such as *Pf*AMA1, *Pf*MSP3, *Pf*MSP7, and *Pf*EBA181) when hepcidin was present (Fig. 2) may suggest an improved ability of the released merozoites to invade erythrocytes. Thus, the addition of the peptide hormone to the culture media could be a signal for the parasite to reduce metabolic processes and to increase its invasion efficiency.

In addition to roles in parasite proliferation and development, different levels of labile iron may induce regulatory processes at various levels. Under high-iron conditions, the observed upregulation of histone deacetylation (Fig. 2C) may lead to the condensation and thus deactivation of certain chromatin regions (88). Similarly, iron-mediated regulation of mRNA translation by iron-regulatory proteins has been described in yeast, trypanosomes and mammals (89–91). The binding sites and target genes of the differentially expressed transcription factors and of *Pf*IRP remain to be identified in *P. falciparum*. Moreover, protein phosphorylation may play a role in iron-dependent regulatory mechanisms. As a serine/threonine kinase (KIN) serves as a nutrient sensor in *P. berghei*, driving a fast response that leads to increased parasite multiplication and virulence (16), a similar kinase may sense iron and lead to increased replication in *P. falciparum*.

On the basis of our RNA-sequencing results (Fig. 2 and Table 1) and the *P. falciparum* transporter list (28), we identified six proteins that are likely involved in iron transport in the parasite (Table 2 and Figure 6) and analyzed their subcellular localization (Fig. 3), their importance for growth (Fig. 4), and their predicted structures (Fig. 5). *Pf*MRS3 transcription was upregulated at the ring stage under high vs. low-iron conditions (log_2_ FC = 0.33, *P* = 0.002, Fig. 2B), and fluorescence of the GFP-tagged protein was exclusively detected at the mitochondrion (Fig. 3A). As a disruption of the gene was reported to fail (65), we were not able to generate a knockdown line after four independent attempts, and parental DNA of the original gene locus was still present in the GFP reporter line (Supplementary Fig. S2), *Pf*MRS3 is likely essential for asexual growth like PBANKA_041620 (E = 1 x 10^-69^, 71.4% identity, 25% coverage) in *P. berghei* (92). The orthologous mitochondrial iron transporter (*Tg*MIT, TGME49_277090, E = 7 x 10^-19^, 26.0% identity, 28% coverage) also localized to the mitochondrion in *T. gondii* and was upregulated at the protein level upon iron overload in consequence of a *Tg*VIT knock out in the related apicomplexan parasite (33). Our structural analyses (Fig. 5, Supplementary Fig. S3A and S4A) further support that *Pf*MRS3 imports ferrous iron into the mitochondrion, the main iron user of the cell, thereby reducing the cytosolic Fe^2+^ concentration (Fig. 6) as a means of detoxification, which has been reported for yeast (93). The protein’s substrate specificity as well as iron binding and transport activity remain to be confirmed experimentally.

**Figure 6:**
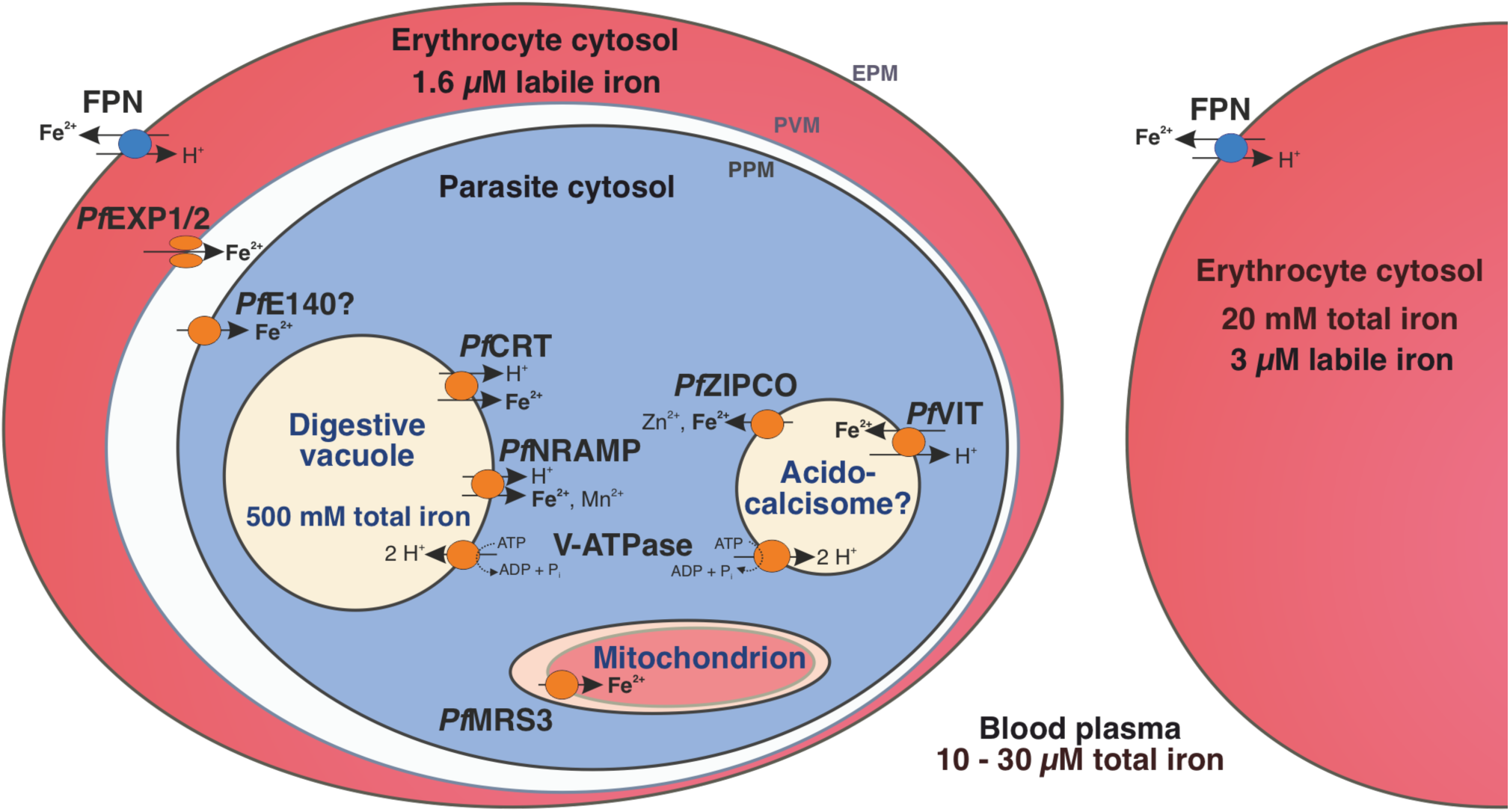
Iron homeostasis in a *P. falciparum*-infected erythrocyte. The human blood plasma contains between 10 and 30 µM total iron and the erythrocyte cytosol approximately 20 mM (19). However, the labile iron pool is only 3 µM in an uninfected erythrocyte and 1.6 µM in a *P. falciparum*-infected one (20). Human ferroportin (FPN) at the host cell surface (erythrocyte plasma membrane, EPM) exports ferrous iron from the erythrocyte (127) and the nutrient pore formed by *Pf*EXP1 and *Pf*EXP2 allows the passage of ions through the parasitophorous vacuole membrane (PVM) (128). *Pf*E140 at the parasite plasma membrane (PPM) may mediate iron uptake into the parasite cytosol and the mitochondrial carrier protein *Pf*MRS3 likely translocates Fe^2+^ into the mitochondrion, a site of *de novo* heme biosynthesis (this study). We propose that the vacuolar iron transporter (*Pf*VIT) is involved in iron detoxification by transporting excess Fe^2+^ from the cytosol into cytoplasmic vesicles that may be acidocalcisomes, whereas *Pf*ZIPCO releases Fe^2+^ from these organelles under low-iron conditions. The digestive vacuole (DV) contains a high amount of total iron as it is the site of hemoglobin degradation and hemozoin formation (21). The chloroquine resistance transporter (*Pf*CRT) and the natural resistance-associated macrophage protein (*Pf*NRAMP, also called *Pf*DMT1 for divalent metal transporter 1) were suggested to mediate proton-coupled export of Fe^2+^ from the DV into the parasite cytosol (37, 73). Both acidocalcisomes and the DV are likely acidified by the plant-like H^+^-pump V-ATPase, which can fuel secondary active transport processes (22, 27). Parasite-encoded proteins are shown in orange and human-encoded transporters in blue.

Complementation assays in *S. cerevisiae* indicated a role for *Pf*VIT in iron detoxification (30, 31) and we observed that the expression of the gene was upregulated under high vs. low-iron conditions in *P. falciparum* (log_2_ FC = 0.29, *P* = 0.02, Fig. 2B). The fluorescence pattern of *Pf*VIT-GFP in live cells (Fig. 3B) was consistent with cytoplasmic vesicles that may be acidocalcisomes, as described for *T. brucei* VIT1 (24). An increase in the number of fluorescent punctate structures during parasite development (Fig. 3B) was also observed for VIT in *T. gondii* (33). *Pf*VIT shares 47.0% identity with *Tg*VIT (E = 8 x 10^-84^, 95% coverage) and 36.9% identity with *Tb*VIT1 (E = 9 x 10^-39^, 98% coverage). In contrast, *P. berghei* VIT (PBANKA_143860, E = 3 x 10^-160^, 79.3% identity, 98% coverage) was shown to localize to the ER in indirect immunofluorescence assays (30). This may be explained by differences between species or variation in methodology such as fixation, permeabilization, and immunolabeling techniques as opposed to live-cell imaging (94–96).

Transport assays using inverted vesicles that were prepared using recombinant *Pf*VIT expressed in *E. coli* demonstrated that the protein is a Fe^2+^/H^+^ antiporter (32). The translocation of Fe^2+^ in exchange for H^+^ is likely fueled by the pH gradient across the membrane of the acidic vesicles and the high similarity of the putative Fe^2+^-binding pocket at the cytosolic side of the predicted *Pf*VIT structure with that of experimentally characterized *Eg*VIT1 (Fig. 5, Supplementary Fig. S4B) provide further evidence for our hypothesis. While not essential during asexual blood stages (65), a knockout of VIT resulted in reduced liver stage development in *P. berghei* (30) and increased sensitivity to high iron levels in both *P. berghei* (30) and *T. gondii* (33). Similarly, growth of the ΔVIT *P. falciparum* line was not affected under standard conditions, whereas the addition of hepcidin – which increases intracellular labile iron levels (Fig. 1A) – compromised parasite proliferation in our study (Fig. 4B). Thus, we hypothesize that the transporter sequesters Fe^2+^ into cytoplasmic vesicles, which is important for iron detoxification under high-iron conditions. While ΔVIT *P. falciparum* is more sensitive to elevated intracellular Fe^2+^ concentrations (Fig. 4B) as a consequence of impaired removal of excess iron from the cytosol, *Pf*MRS3 may compensate for a loss of *Pf*VIT under standard conditions by transporting ferrous iron from the cytosol into the mitochondrion (Fig. 6).

In contrast to the PPM staining of *P. berghei* sporozoites in immunofluorescence assays, *Pf*ZIPCO-GFP expression resulted in a punctate fluorescence pattern in the cytoplasm of live blood-stage *P. falciparum* (Fig. 3C), similar to that of *Pf*VIT-GFP (Fig. 3B). Whereas the *Pf*ZIPCO knockout caused a growth increase under standard conditions (Fig. 4B), ΔZIPCO *P. berghei* parasites displayed normal blood-stage development but impaired sporozoite infectivity as well as reduced replication at the liver stage in mice (34). Interestingly, the ortholog TGME49_225530 is also dispensable in *T. gondii* tachyzoites with a phenotype score of −2.94 (values below −1.5 are considered non-essential (97)). Hence, Fe^2+^ efflux from cytoplasmic vesicles (potentially acidocalcisomes) into the cytosol via *Pf*ZIPCO may be dispensable in *P. falciparum* under iron-replete conditions during the blood stage because of the redundancy with iron import mechanisms into the parasite, and the production of the protein may result in a fitness cost. In contrast, liver-stage parasites in low-iron environments may rely on the transporter’s activity when the demand for iron is high during schizogony. As the transcription of *Pf*ZIPCO was upregulated at low vs. control iron levels (log_2_ FC = 0.55, *P* = 0.04, Table 1) and in response to hepcidin treatment (log_2_ FC = 0.63, *P* = 0.006, Fig. 2D, Table 1), the transport protein may release Fe^2+^ and Zn^2+^ ions from intracellular stores, in this case cytoplasmic vesicles (Fig. 6), in case of scarcity, thereby increasing cytosolic ion levels like other ZIP transporters (43). While our analyses of the predicted structure and its alignment with *Bb*ZIP indicate that *Pf*ZIPCO likely has the capacity to bind and transport cations like Fe^2+^ or Zn^2+^ (Fig. 5, Supplementary Fig. S3C and S4C), its substrate specificity can only be conclusively established by characterizing the purified protein. Liposomal assays with the putative zinc transporter *Pf*ZIP1 (PF3D7_0609100, 24.5% identity with *Pf*ZIPCO, E = 1 x 10^-19^, 78% coverage), which localized to the plasma membrane in schizonts, demonstrated that this ZIP transporter preferentially binds Zn^2+^ over Fe^2+^ (98). Interestingly, the preference was abolished if the histidine-rich loop at the C-terminus of *Pf*ZIP1, which is not present in *Pf*ZIPCO, was truncated. As mRNA levels of *Pf*ZIP1 were enhanced at low cytosolic Zn^2+^ levels (98) but not differentially regulated under various iron conditions (Supplementary Tables S1 and S2), it may play a role in zinc rather than iron homeostasis under physiological conditions.

As the highest intracellular iron concentration in *P. falciparum* is reached within the DV (21, 99), the free form of the metal may need to be exported from this compartment under high-iron conditions to prevent damage to the DV membrane (Fig. 6). This function may be fulfilled by *Pf*CRT (37) and / or *Pf*NRAMP (73), which were both upregulated under high vs. low-iron conditions in our RNA-sequencing analysis (log_2_ FC = 0.26, *P* = 0.007 and log_2_ FC = 0.28, *P* = 0.003, respectively, Fig. 2B, Table 1) and are essential in asexual parasites (35, 36, 65). The predicted structure of *Pf*NRAMP (Fig. 5) reflects the state that is open towards the cytosol as in the crystal structure of NRAMP from *Deinococcus radiodurans* (100). While a negatively charged cavity inside the protein is clearly visible in the *Pf*NRAMP model, the proposed outward-facing permeation pathway for metal ions is likely occluded in this conformation (Fig. 5C). It is conceivable that Fe^2+^ ions permeate through this pathway from the DV lumen and bind to the charged cavity like the Mn^2+^ ion to *S. capitis* NRAMP/DMT (Supplementary Fig. S4D). *Pf*NRAMP might function similarly to its ortholog in *D. radiodurans*, which was shown to mediate pH-dependent transport of Fe^2+^ and Mn^2+^ in symport with H^+^ using uptake assays in *E. coli*, HEK293T cells, and proteoliposomes (100, 101).

Expression of the surface protein *Pf*E140 was upregulated when iron levels were low compared to standard conditions (log_2_ FC = 0.65, *P* = 0.0006, Table 1) and the GFP fusion protein localized to the PPM only, as evidenced by the fluorescent edges of free merozoites (Fig. 3D). This observation is consistent with the fact that the extracellular portions of this protein are highly polymorphic because of their exposure to the immune system at the sporozoite stage (76). Interestingly, vaccines targeting *Py*E140 in *Plasmodium yoelii* were reported to induce up to 100% sterile protection mediated by antibodies in mice (102). The reduced parasite replication rate upon its conditional knockdown demonstrates the importance of *Pf*E140 for parasite growth and the rescue of the *Pf*E140 knockdown by hepcidin treatment support a role of this putative transporter in iron uptake (Fig. 4F). Its predicted essential nature (65), in addition to the absence of orthologs in humans, make it an excellent drug target candidate. While our *P. falciparum* gene expression data (Fig. 2B and D) point towards a role of *Pf*E140 in iron homeostasis, its precise function is still unclear and it remains to be clarified whether the large coiled-coil domain exposed to the extracellular space (Fig. 5) can mediate dimerization upon substrate binding. Given our experimental results and the functional annotations (Fig. 5C, Supplementary Fig. S5), we hypothesize that *Pf*E140 is a plasma membrane transporter for inorganic cations such as metal ions.

In conclusion, this is the first study to investigate *P. falciparum* transcriptomics under different iron conditions and to determine the subcellular localization of the known and putative iron transport proteins *Pf*MRS3, *Pf*VIT, *Pf*ZIPCO and *Pf*E140 as well as the growth effects of a *Pf*VIT or *Pf*ZIPCO knockout and an inducible *Pf*E140 knockdown. Our results reveal how the human malaria parasite reacts to alterations in host iron status and provide new insights into the mechanisms of iron transport in *P. falciparum* in addition to offering avenues for the development of novel therapeutic strategies against malaria. We propose a model for the regulation of iron homeostasis in the *P. falciparum*-infected erythrocyte with a series of six organelle-specific iron transport proteins in the parasite (Fig. 6): One route of iron uptake into the parasite is through the release of Fe^2+^ upon hemoglobin digestion in the DV and the efflux of the ion into the cytosol mediated by *Pf*NRAMP (38) and / or *Pf*CRT (37). Ferrous iron likely also enters the parasite cytosol across the PPM via *Pf*E140 and this pathway may be particularly important during schizogony, when the putative transporter gene is abundantly transcribed and new merozoites without a DV are formed (51). Once inside the cytosol, iron concentrations need to be tightly regulated to avoid toxicity, which could be achieved by Fe^2+^ import into the mitochondrion as the main site of iron utilization via *Pf*MRS3 and through transport into (*Pf*VIT) and out of (*Pf*ZIPCO) cytoplasmic vesicles functioning as labile iron pools. To confirm the hypotheses of our exploratory study, transport assays with purified proteins like those performed with recombinant *Pf*VIT (32) are required for the formal demonstration of substrate specificities and activities of the other transporters in addition to further functional characterization of the proteins during the mosquito, liver and asexual blood stages of the parasite. As no ortholog of the essential *Pf*E140 and only a distant homolog of the non-redundant mitochondrial iron importer *Pf*MRS3 (42) are present in humans, these provide candidate targets for urgently needed new antimalarial drugs. Furthermore, dissecting how *P. falciparum* senses changes in micronutrient availability in its environment and how it modulates its virulence accordingly is an area of considerable interest for future investigation, as iron is an essential regulatory signal for virulence factors in many pathogens.

## MATERIALS AND METHODS

### *P. falciparum* culture and proliferation assays

The *P. falciparum* strain 3D7 was cultured according to modified standard procedures (103) at 5% hematocrit using human 0 Rh+ erythrocytes from the University Medical Center Hamburg-Eppendorf (UKE), Germany, at 1% O_2_, 5% CO_2_ and 94% N_2_. RPMI 1640 medium was supplemented with 0.5% (w/v) AlbuMAX II, 20 µg/mL gentamicin and 100 µM hypoxanthine (Thermo Fisher Scientific). Mature schizonts were obtained by treating schizonts at 40 hpi with 1 mM compound 2 (4-[7-[(dimethylamino)methyl]-2-(4-fluorphenyl)imidazo[1,2-α]pyridine-3-yl]pyrimidin-2-amine, LifeArc) for 8 h. To count the number of merozoites per mature schizont, Giemsa-stained blood smears were analyzed by light microscopy. Only single-infected cells with one digestive vacuole were taken into account.

To assess parasite proliferation over six days, a previously described assay on the basis of flow cytometry was employed (104). Parasites were synchronized to a 3-h age window by isolating late schizonts from a 60% Percoll (GE Healthcare) gradient and culturing these for 3 h with fresh erythrocytes (105), followed by controlled elimination of advanced parasite stages using 5% (w/v) D-sorbitol (Carl Roth) for 10 min at 37°C (106). The growth assay was started at 0.1% parasitemia using the resulting ring-stage parasites at 0 – 3 hpi. The parasitemia was determined at the trophozoite stage every two days by flow cytometry and culture media with the respective supplements were exchanged daily.

### Flow cytometry

To determine parasitemia, 20 µL of resuspended parasite culture was added to 80 μL culture medium and stained with 5 μg/mL SYBR Green I (Thermo Fisher Scientific) and 4.5 μg/mL dihydroethidium (DHE, Sigma-Aldrich) in the dark for 20 min at room temperature. Stained cells were washed with PBS three times and analyzed with an ACEA NovoCyte flow cytometer and NovoExpress Software (version 1.6.1, Agilent). Forward and side scatter gating was used to identify erythrocytes and SYBR Green I fluorescence intensity to determine the number of parasitized cells per 100,000 events recorded for each replicate. For Phen Green SK measurements, uninfected erythrocytes were washed with PBS and incubated with 10 µM Phen Green SK in PBS at 37°C for 60 min. DHE at 4.5 μg/mL was added during the last 20 min of incubation. After three washes with PBS, the cells were analyzed as described above.

### Cloning of DNA constructs

For generating the GFP reporter lines, a homologous region of approximately 800 bp at the 3’ end of the respective gene was amplified without the stop codon from 3D7 gDNA using Phusion high fidelity DNA polymerase (New England Biolabs). A homology region of about 400 bp at the 5’ end of the respective gene was used for targeted gene disruption. The fragments were then inserted into pSLI-GFP (107) using Not*I* and Avr*II* restriction sites. For *glmS* constructs, pSLI-GFP-*glmS* (108) was used as a vector instead. All oligonucleotides and plasmids used in this study are listed in Supplementary Table S3.

### Transfection of *P. falciparum*

As described previously (109), parasites at the late schizont stage were purified using 60% Percoll (105) and electroporated with 50 μg DNA of the respective plasmid in a 0.2-cm gap cuvette (Bio-Rad Laboratories) using Amaxa Nucleofector 2b (Lonza). Either 4 nM WR99210 (Jacobus Pharmaceuticals) or 2 μg/mL blasticidin S (Life Technologies) was used for selecting transfectants. For the selection of parasites that were genomically modified using the SLI system (107), 400 μg/mL G418 (ThermoFisher Scientific) was added to the culture medium once the parasitemia reached 5%. After the selection of modified parasites, genomic DNA was isolated with the QIAamp DNA Mini Kit (Qiagen) and diagnostic tests for correct integration into the genome were performed as specified earlier (107).

### Confocal live-cell microscopy

Erythrocytes infected with parasites at different stages at 3 – 6% parasitemia were incubated in culture medium with 20 nM MitoTracker Red, 200 nM ER Tracker Red or 100 nM LysoTracker Deep Red (Invitrogen, if applicable) at 37°C for 20 min. Then, 200 nM Hoechst-33342 (Invitrogen) was added for 10 min prior to washing the cells with Ringer’s solution (122.5 mM NaCl, 5.4 mM KCl, 1.2 mM CaCl_2_, 0.8 mM MgCl_2_, 11 mM D-glucose, 25 mM HEPES, 1 mM NaH_2_PO_4_, pH 7.4) prewarmed to 37°C and seeding on a chambered No. 1.5 polymer cover slip (Ibidi). After 5 min, unbound erythrocytes were removed by washing with Ringer’s solution and the sample was placed into an incubation chamber that maintained the microscope work area including the objective at 37°C. Images and videos were acquired using an SP8 confocal microscope system with a 63x oil-corrected lens (C-Apochromat, numerical aperture = 1.4) and Lightning deconvolution software (Leica), and processed using ImageJ version 2.9.0/1.53t (110). If fluorescence intensities were to be quantified, no averaging or deconvolution software was applied.

### Conditional knockdown mediated by *glmS* ribozyme

For *glmS*-based knockdown induction (66), highly synchronous parasites at the early ring stage were cultured with or without supplementation with 2.5 mM glucosamine (GlcN, Sigma-Aldrich). The knockdown was quantified by confocal live-cell microscopy using schizonts 36 h post GlcN treatment initiation. Images of parasites of similar size were acquired with the same settings and background-corrected fluorescence intensities (integrated density) as well as the size of the region of interest were determined using ImageJ version 2.9.0/1.53t (110), and the data visualized using Graph Pad Prism version 9.4.1.

### Sample collection for RNA extraction and RNA sequencing

Parasites were synchronized to a three-hour window after invasion of erythrocytes from the respective donor as described above. Samples for RNA-sequencing were prepared in triplicate for each condition and time point, i.e., three separate parasite cultures each were grown in parallel for a total of at least two weeks. During the second IDC, two 10-mL dishes each were harvested for parasites at the ring stage (6 – 9 hpi) and one 10-mL dish each for trophozoites (26 – 29 hpi). Samples were collected by centrifuging the culture for 5 min at 800 g and 37°C and dissolving the erythrocyte pellet using 5 mL TRIzol (ThermoFisher Scientific) prewarmed to 37°C, followed by immediate transfer to −80°C for storage. The parasitemia was 0.3% at the start of the experiments with high, control and low-iron donor blood and 2 – 3% at the time of harvest. Parasite cultures treated with 0.7 µM hepcidin (Bachem) had a starting parasitemia of 0.6% and untreated cultures 1% to reach a parasitemia of 4 – 5% during the second cycle. For each experiment, the parasitemia was kept consistent at the point of harvest as high parasite densities can affect transcription (15).

For RNA extraction, the samples frozen in TRIzol were thawed, mixed thoroughly with 0.1 volume cold chloroform, and incubated at room temperature for 3 min. Following centrifugation at 20,000 g and 4°C for 30 min, the supernatants were transferred to fresh vials and combined with 70% ethanol of equal volume. RNA was purified using the RNeasy MinElute Kit (Qiagen) by on-column DNase I digest for 30 min and elution with 14 µL water. The GLOBINclear Human Kit (ThermoFisher Scientific) was then employed to deplete human globin mRNA in all samples. The Qubit RNA HS Assay Kit and Qubit 3.0 fluorometer (ThermoFisher Scientific) were used for RNA quantification. Upon arrival at the EMBL Genomics Core Facility (GeneCore Heidelberg, Germany), the RNA quality of each sample was evaluated using the RNA 6000 Nano kit and Bioanalyzer 2100 (Agilent). The median RNA integrity number (RIN) of all samples was 7.30 (IQR: 6.85 – 8.15, Supplementary Fig. S1). Individually barcoded strand-specific libraries for mRNA sequencing were prepared from total RNA samples of high quality (approximately 150 ng per sample) using the NEBNext® RNA Ultra II Directional RNA Library Prep Kit (New England Biolabs) for 12 PCR cycles on the liquid handler Biomek i7 (Beckman Coulter) at GeneCore. Libraries that passed quality control were pooled in equimolar amounts, and a 2 pM solution of this pool was sequenced unidirectionally on a NextSeq® 500 System (Illumina) at GeneCore, resulting in about 500 million reads of 85 bases each.

### RNA sequencing read mapping and data analysis

Following successful initial quality control of the RNA-sequencing reads with FastQC version 0.11.8 (111), sequencing adapters were trimmed using Cutadapt version 2.10 (112). A genome index was generated using the FASTA sequence file of the *P. falciparum* 3D7 genome release 46 (PlasmoDB-46_Pfalciparum3D7_Genome.fasta) and the GFF3 annotation file (PlasmoDB-46_Pfalciparum3D7.gff), both obtained from PlasmoDB (113), with STAR version 2.7.5c (114). The same R package was used to align reads to the genome with a maximum of three allowed mismatches (--outFilterMismatchNmax 3). To consolidate the results obtained with FastQC and STAR alignments, a single report file was created using MultiQC version 1.9 (115).

The mapped reads were then summarized in Sequence Alignment/Map (SAM) format using featureCounts (116) from the R package Rsubread version 2.2.1 (117). For counting mapped reads per gene using featureCounts, fragments with a minimum length of 50 bases were considered (minFragLength = 50). Therefore, gene IDs and lengths of transcripts were extracted from PlasmoDB-46_Pfalciparum3D7_AnnotatedTranscripts.fasta with SAMtools faidx version 1.10.2 (118). The R package edgeR 3.30.3 (119) was used to compute RPKM values (reads per kilobase per million mapped reads) and for differential gene expression analysis. Gene annotations were retrieved from PlasmoDB (113) and PhenoPlasm (120). The results of these analyses were visualized with volcano plots using the R package Enhanced Volcano version 1.15.0 (121). The raw and processed data (FASTA files, RPKM values and results of the differential gene expression analysis) can be accessed at https://www.ebi.ac.uk/biostudies/studies/E-MTAB-13411.

The highly polymorphic *var*, *stevor*, and *rifin* gene families were excluded from downstream analyses because of their great sequence diversity between parasites of the same strain during mitotic growth (58, 59). Genes that were significantly regulated (defined as *P* < 0.05 according to the exact test for the negative binomial distribution with Benjamini-Hochberg correction (54) and an absolute value of log_2_ FC ≥ 0.2) were subjected to functional enrichment analysis with g:Profiler (https://biit.cs.ut.ee/gprofiler/gost (60), accessed on August 17, 2022). The resulting GO, KEGG and REAC terms were summarized using REVIGO (http://revigo.irb.hr/) with the similarity value set to 0.5 (122) and visualized as in Thomson-Luque et al. (83) using the scientific color map “roma” (123). To estimate parasite age, an algorithm developed by Avi Feller and Jacob Lemieux (50) was adapted to use expression data from Broadbent et al. (51) with the time points 6, 14, 20, 24, 28, 32, 36, 40, 44, and 48 hpi as reference. The code and data used were deposited to Zenodo with the record ID 7996302 (https://zenodo.org/record/7996302).

### Protein structure prediction

Structure predictions for monomeric proteins were obtained from AlphaFold Protein Structure Database version 3 (67, 68) and homodimeric proteins were predicted using AlphaFold2-multimer version 2.2.2, database version 2.2.0 (69) deployed at the EMBL Hamburg computer cluster. Molecular visualization was performed with UCSF ChimeraX version 1.3 (124). UCSF Chimera MatchMaker and Match → Align tools with default settings were used for structural comparison of the predicted structures of *P. falciparum* proteins with putative orthologs and sequence alignments were generated using the Match → Align tool (125). The DeepFRI server (https://beta.deepfri.flatironinstitute.org) was used to identify possible functional residues with the DeepFRI graph convolutional network (77).

## Supporting information

PfVIT.pdb

PfZIPCO.pdb

## Supplementary Material

### Supplementary Figures

**Supplementary Figure S1:** RNA quality.

**Supplementary Figure S2:** Cloning strategy and confirmation of correct DNA integration into the genome of the cell lines generated.

**Supplementary Figure S3:** Sequence alignments of *P. falciparum* proteins with functionally characterized homologs.

**Supplementary Figure S4:** Alignments of functional sites in the predicted *P. falciparum* protein structures with those of functionally characterized homologs.

### Supplementary Videos

**Supplementary Video S1:** Parasite endogenously expressing *Pf*MRS3-GFP stained with MitoTracker Red and Hoechst-33342.

**Supplementary Video S2:** Parasite endogenously expressing *Pf*VIT-GFP stained with ER Tracker Red and Hoechst-33342.

**Supplementary Video S3:** Parasite endogenously expressing *Pf*ZIPCO-GFP stained with LysoTracker Deep Red and Hoechst-33342.

**Supplementary Video S4:** Parasite expressing *Pf*E140-GFP endogenously, *Pf*ACP(1–60)-mCherry episomally and stained with Hoechst-33342.

### Supplementary Tables

**Supplementary Table S1:** Differentially expressed genes of *P. falciparum* 3D7 cultured with erythrocytes from a donor with higher, control or low iron status.

**Supplementary Table S2:** Differentially expressed genes of *P. falciparum* 3D7 cultured with erythrocytes from a healthy donor with or without addition of 0.7 µM hepcidin.

**Supplementary Table S3:** Oligonucleotides (A) and plasmids (B) used in this study.

### Supplementary PDB files

Top-scoring PDB files for the multimers that were computed at EMBL:

**PfVIT.pdb**

**PfZIPCO.pdb**

## Acknowledgements

The authors thank the Genomics Core Facility at EMBL Heidelberg, especially Vladimir Benes, for the RNA-sequencing service, and EMBL Hamburg for the provision of research and technical support as well as access to research infrastructures. Grzegorz Chojnowski and the group of Jan Kosinski at EMBL Hamburg enabled the AlphaFold2 workflow at the EMBL Hamburg computer cluster. The Advanced Light and Fluorescence Microscopy Facility at CSSB Hamburg, in particular Roland Thünauer, supported microscopy experiments and the Bernhard Nocht Institute for Tropical Medicine (BNITM) provided lab space. We gratefully acknowledge Tobias Spielmann for pSLI-GFP and pARL-*Pf*ACP(1–60)-mCherry, Paul Burda for pSLI-GFP-*glmS*, Jacobus Pharmaceuticals for WR99210, Anna Bachmann and Mayka Sánchez for helpful advice, Eileen Devaney, Katharina Jungnickel, and Samuel Pažický for critical reading of the manuscript, and Heidrun von Thien, Yannick Höppner, and Gabriela Guédez for technical assistance.

## Author contributions

JS and JW designed the study; MG and SPe recruited the blood donors; JW and CN performed the experiments; JW, JS, VK and LVN analyzed the data; VK generated the structural models, and JW wrote the manuscript with contributions from VK, JS and SPo. All authors read and approved the submitted version.

