## Supplementary Figures for "Iron transport pathways in the human malaria parasite *Plasmodium falciparum* revealed by RNA-sequencing"

**Experiment 1:**

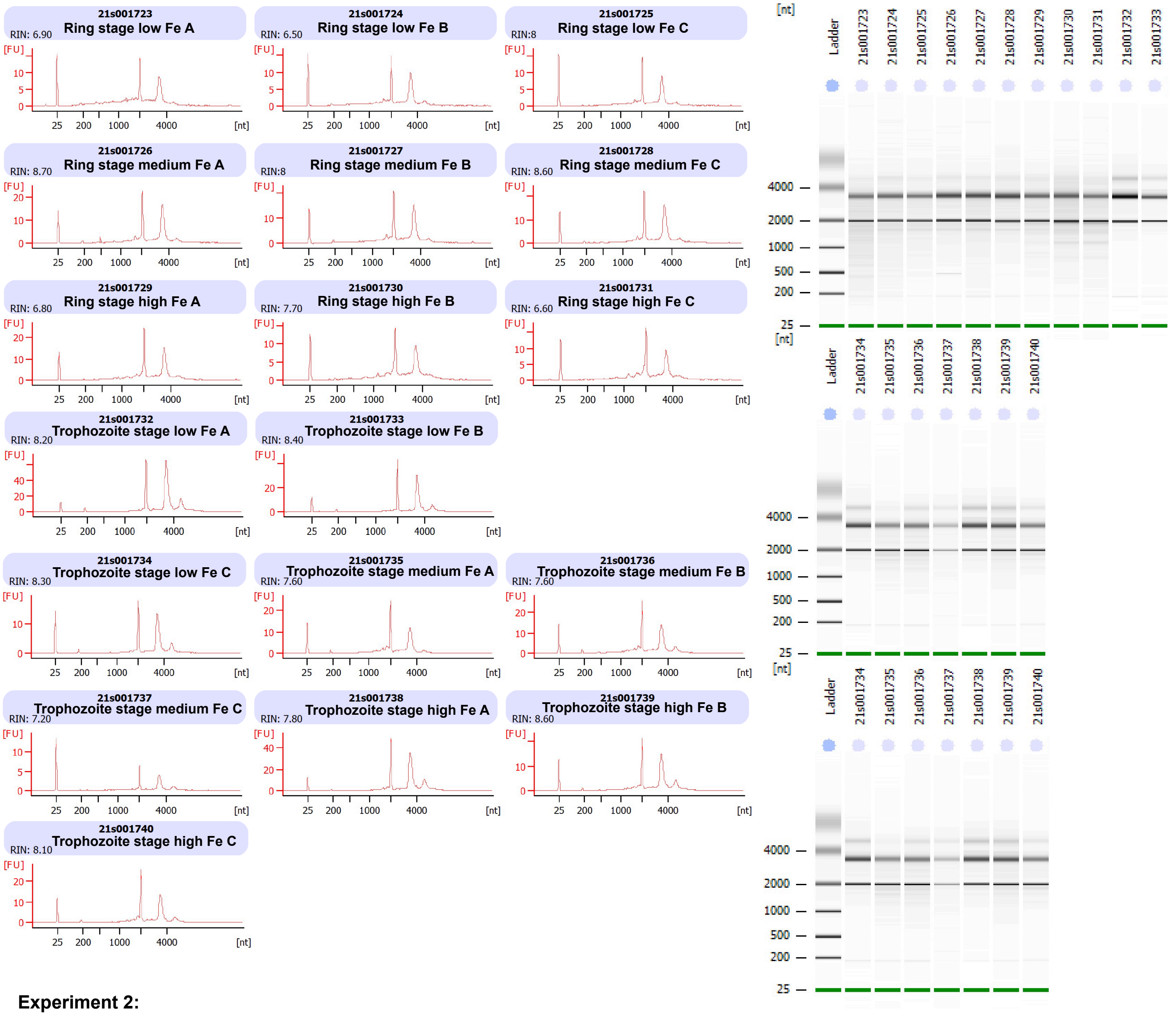


**Experiment 2:**

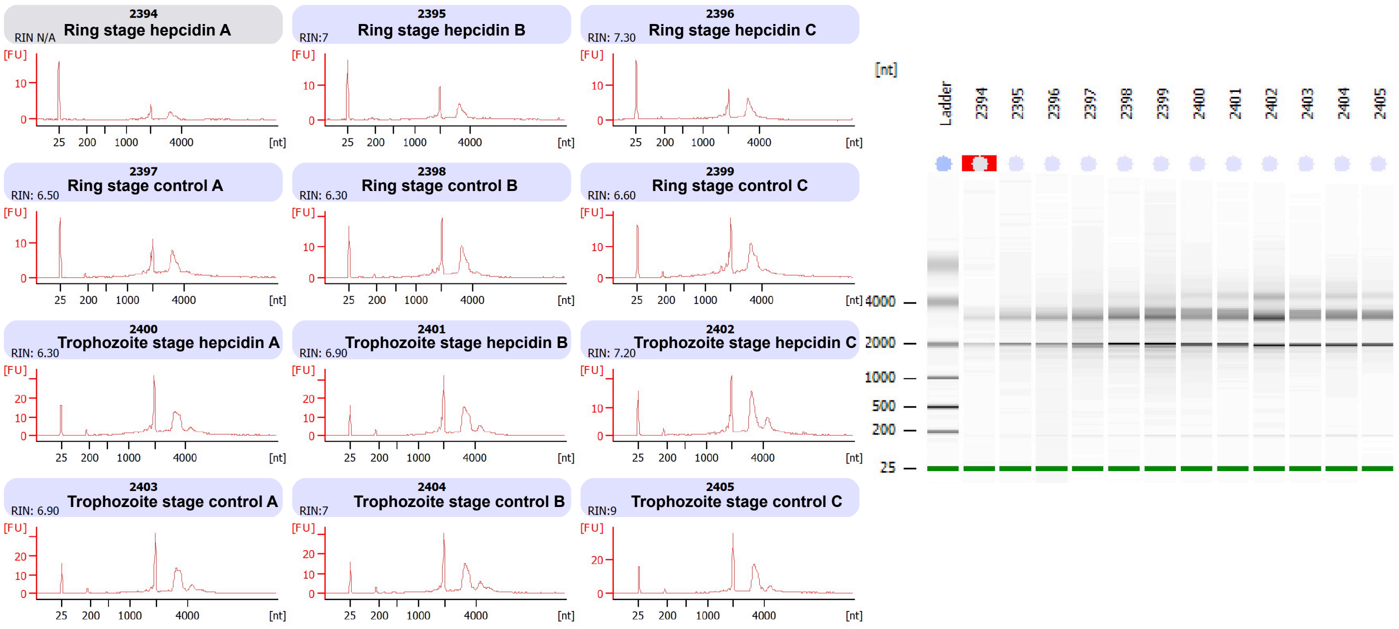


**Supplementary Figure S1: RNA quality.**
Before library synthesis, the quality of total RNA was assessed for all samples using Bioanalyzer 2100 (Agilent). The RNA integrity number (RIN) is indicated and the peaks at approximately 2,000 and 3,600 nucleotides (nt) correspond to *Plasmodium falciparum* 18S and 28S rRNA. FU, fluorescence units.

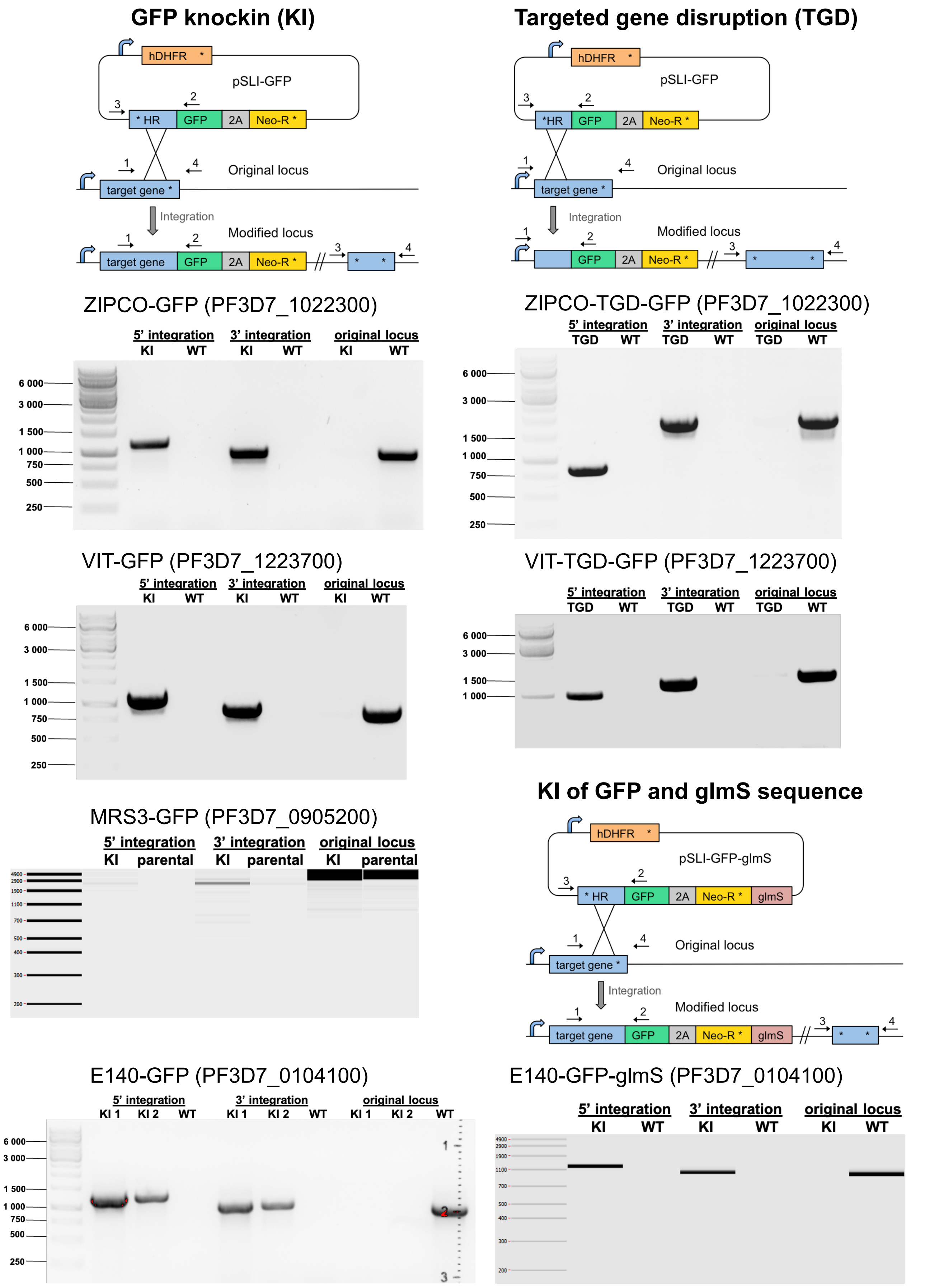


**Supplementary Figure S2: Cloning strategy and confirmation of correct DNA integration into the genome of the cell lines generated.**
Schematics depict the endogenous tagging strategy using the selection-linked integration system (SLI). GFP, green fluorescent protein; glmS, glmS ribozyme sequence; hDHFR, human dihydrofolate reductase gene; HR, homology region; Neo-R, neomycin resistance gene; WT, wild-type; 2A, T2A skip peptide. Asterisks represent stop codons and arrows primers (1 to 4) used for diagnostic PCRs.

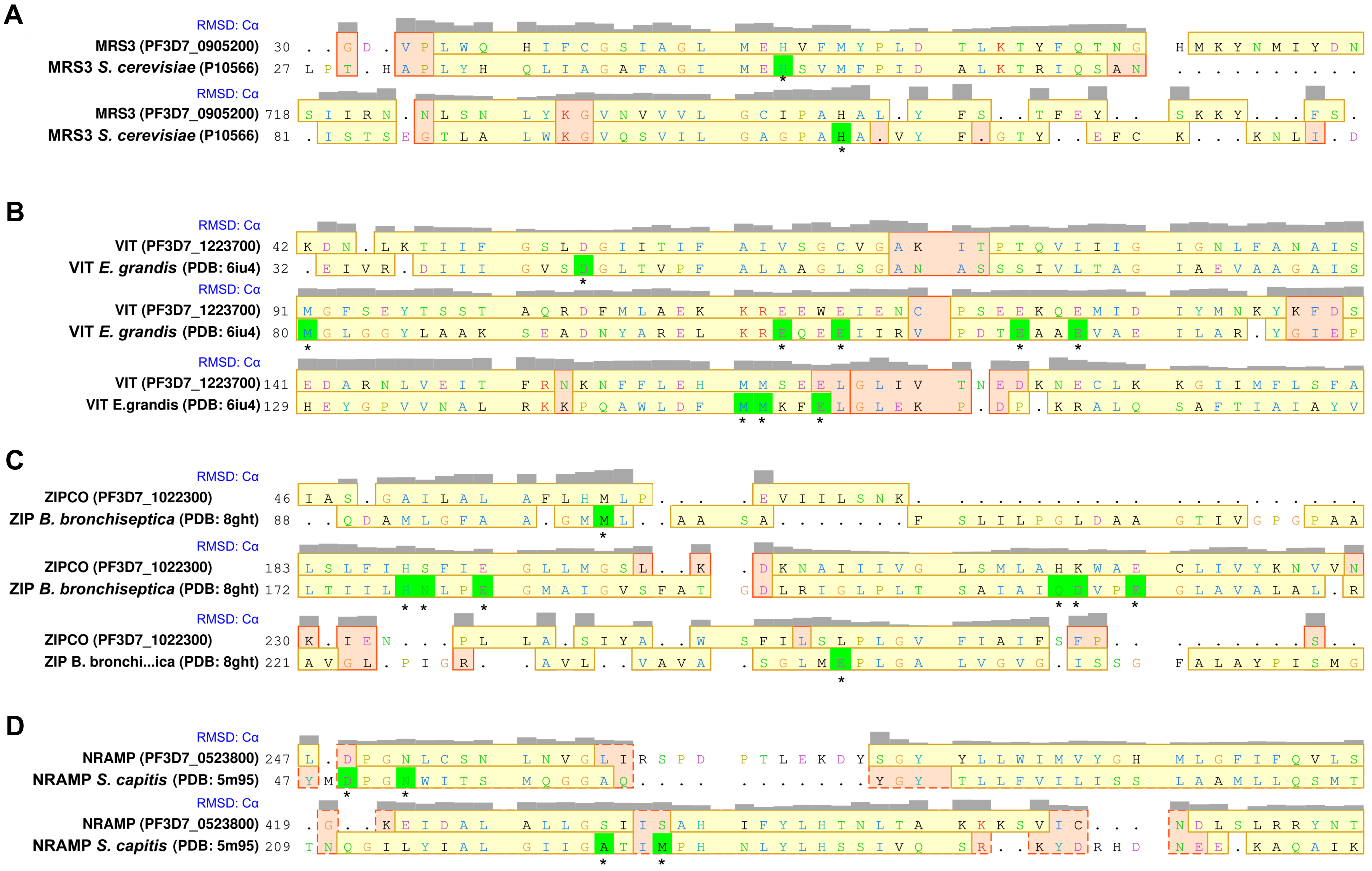


**Supplementary Figure S3: Sequence alignments of *P. falciparum* proteins with functionally characterized homologs.**
The alignments were built by comparison of protein structures using UCSF Chimera MatchMaker and the Match → Align tool. Gray bars indicate Cα RMSD after superposition; the maximum height of the bar corresponds to 5 Å and gaps are indicated as dots. Amino acid residues were colored according to the ClustalW scheme (https://www.jalview.org/help/html/colourSchemes/clustal.html). Yellow background of the sequence stretch corresponds to α-helices and red background stands for an agreement of the sequence that has no secondary structure. Known functional residues that are important for cation binding and/or transport are highlighted with asterisks and a bright green background of the sequence stretch.

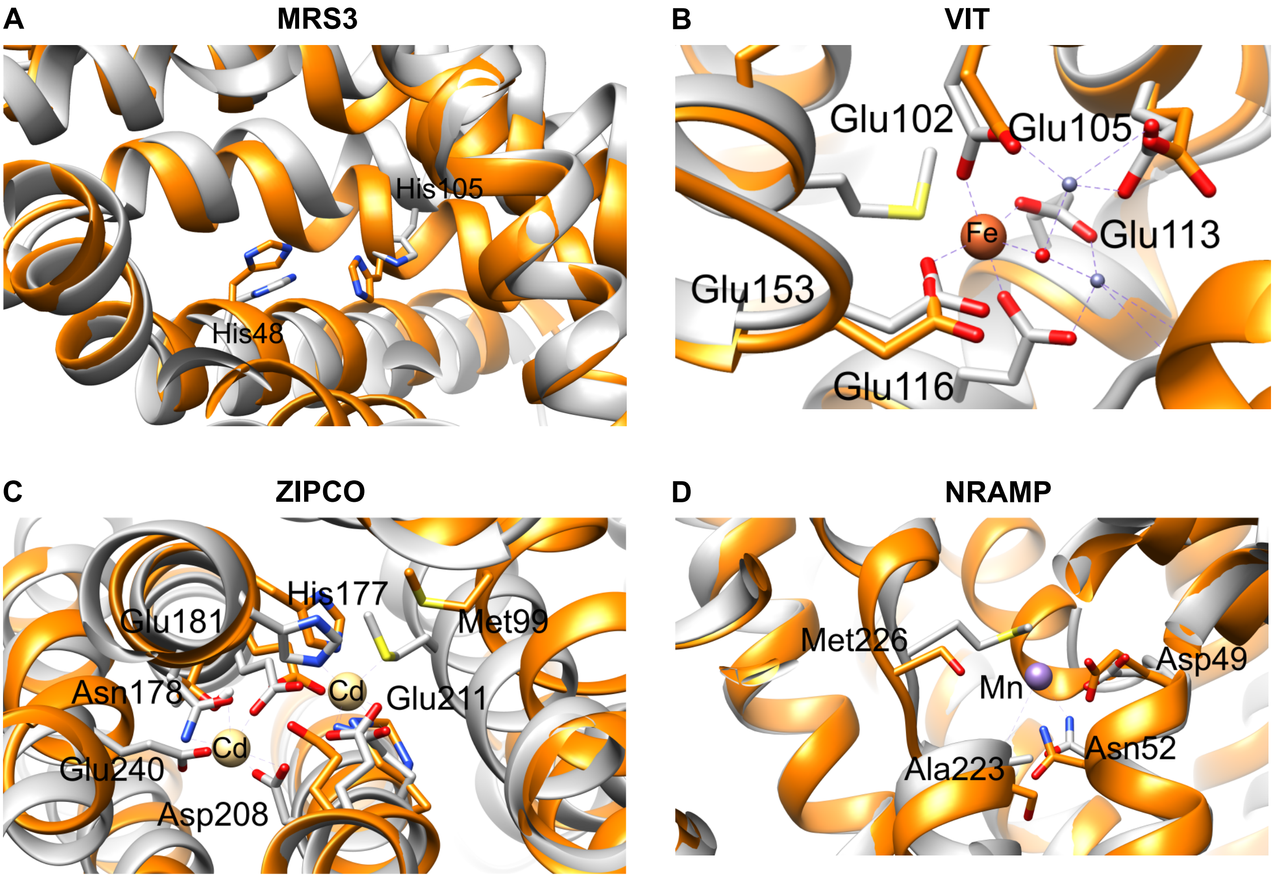


**Supplementary Figure S4: Alignments of functional sites in the predicted *P. falciparum* protein structures with those of functionally characterized homologs.**
Structural alignments were generated using UCSF Chimera MatchMaker and Match → Align tools with default settings. AlphaFold2 predictions for the indicated *P. falciparum* proteins are shown in orange and the structures of their homologs in gray. With the exception of *Saccharomyces cerevisiae* MRS3 (AlphaFold2 prediction P10566), the structures of the homologs were obtained experimentally: VIT1 from *Eucalyptus grandis* (PDB: 6IU9), ZIP from *Bordetella bronchiseptica* (PDB: 8GHT) and NRAMP/DMT from *Staphylococcus capitis* (PDB: 5M95). Key residues and bound ions in the structures of the characterized homologs are indicated and heteroatoms are colored according the conventions.

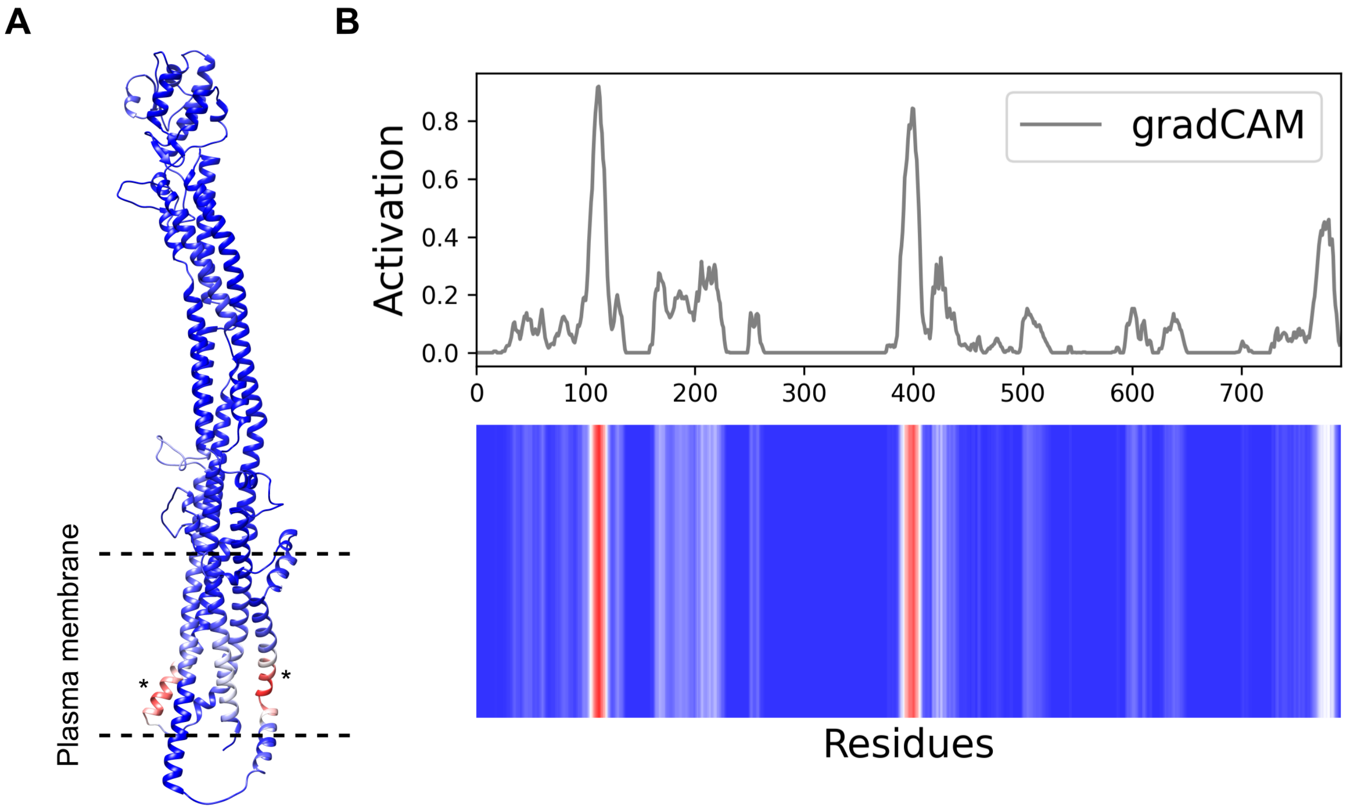


**Supplementary Figure S5: Identification of functional residues in *Pf*E140 using DeepFRI.** **A** The predicted structure of *Pf*E140 is colored according to DeepFRI gradCAM scores for the functional term GO:0015075 “monoatomic ion transmembrane transporter activity” with blue indicating a score of 0 (low confidence) and red indicating a score of 1 (high confidence). The regions of high confidence are located in the transmembrane domains and are indicated with asterisks. **B** Per-residue DeepFRI gradCAM scores for GO:0015075 in *Pf*E140.
